# Leveraging a surrogate outcome to improve inference on a partially missing target outcome

**DOI:** 10.1101/2020.11.29.403063

**Authors:** Zachary R. McCaw, Sheila M. Gaynor, Ryan Sun, Xihong Lin

## Abstract

Sample sizes vary substantially across tissues in the Genotype-Tissue Expression (GTEx) project, where considerably fewer samples are available from certain inaccessible tissues, such as the substantia nigra (SSN), than from accessible tissues, such as blood. This severely limits power for identifying tissue-specific expression quantitative trait loci (eQTL) in undersampled tissues. Here we propose Surrogate Phenotype Regression Analysis (Spray) for leveraging information from a correlated surrogate outcome (e.g. expression in blood) to improve inference on a partially missing target outcome (e.g. expression in SSN). Rather than regarding the surrogate outcome as a proxy for the target outcome, Spray jointly models the target and surrogate outcomes within a bivariate regression framework. Unobserved values of either outcome are treated as missing data. We describe and implement an expectation conditional maximization algorithm for performing estimation in the presence of bilateral outcome missingness. Spray estimates the same association parameter estimated by standard eQTL mapping and controls the type I error even when the target and surrogate outcomes are truly uncorrelated. We demonstrate analytically and empirically, using simulations and GTEx data, that in comparison with marginally modeling the target outcome, jointly modeling the target and surrogate outcomes increases estimation precision and improves power.

## 1 Introduction

Tissue-specific expression quantitative trait loci (eQTL) are of substantial biological interest as mechanisms for explaining how the genetic variants identified in genomewide association studies (GWAS) influence complex traits and diseases [1, 2, 3, 4, 5]. Traditional eQTL studies have focused on accessible tissues such as blood [6, 7], while eQTL discovery in inaccessible tissues, such as the substantia nigra (SSN), have been impeded by insufficient sample sizes. Cross-tissue studies, including the Genotype-Tissue Expression Project (GTEx), have demonstrated that the effect sizes of eQTL are heterogeneous across tissues [8]. Consequently, studying only accessible tissues is insufficient to understand the genetic basis of gene regulation. Larger sample sizes are needed to provide sufficient power for reliable eQTL detection in inaccessible tissues, and there is great interest in borrowing information from accessible tissues to increase the effective sample sizes of inaccessible tissues.

Our work was motivated by the goal of improving power for eQTL mapping in the SSN, a region of the midbrain implicated in the development of Parkinson’s disease [9]. Due to the scarcity of expression data, no previous studies have focused on eQTL mapping in this region. At the time of our analysis, only 80 genotyped subjects with expression data in SSN were available from GTEx, in contrast to 369 with expression in whole blood. Among subjects with expression in blood, nearly 90% were missing expression in SSN. The methodology developed here leverages gene expression from a correlated surrogate tissue, such as blood, to improve power for identifying eQTL in the target tissue, SSN.

Several methods have been developed to address the related problem of multi-tissue eQTL mapping. [10] developed eQtlBma, a fixed-effects, heteroscedastic ANOVA model that jointly models gene expression in multiple tissues. Evidence against the global null hypothesis, that a SNP has no effect on gene expression in any tissue, is quantified using a Bayes factor averaged across potential non-null configurations. [11] proposed Meta-Tissue, which jointly estimates the effect of a SNP on gene expression in multiple tissues using a mixed-effects model, then combines effect size estimates across tissues via meta-analysis. [12] developed MT-eQTL and its extension HT-eQTL, which modeles the vector of Fisher-transformed genotype-expression correlations across tissues. They propose a generative hierarchical model for the multivariate correlation vector and an empirical Bayes procedure for identifying multi-tissue eQTL based on the local false discovery rate.

Our approach differs from existing methods in two key respects. First, we are interested in identifying target-tissue eQTL not multi-tissue eQTL. That is, our null hypothesis is that a SNP has no effect on gene expression in the target tissue, not that a SNP has no effect on gene expression in any tissue. Moreover, we focus on the setting where the target tissue is subject to missing data, and empower eQTL analysis of the target tissue by leveraging data from the surrogate. Second, we are interested in frequentist rather than Bayesian inference, and specifically in asymptotic inference, which does not depend on computationally-intensive permutation procedures that are intractable at genome-scale.

In this paper, we propose improving power for eQTL mapping in an inaccessible tissue (e.g. SSN), for which expression is partially missing, by augmenting the sample with expression data from an accessible surrogate tissue, for which the sample size is substantially larger. Specifically, we propose jointly modeling expression in the target and surrogate tissues while regarding unobserved measurements in either tissue as missing data. We refer to this approach as Surrogate Phenotype Regression Analysis (Spray). Spray leverages the correlation in expression levels across tissues to increase the effective sample size, but maintains eQTL in the target tissue as the focus of inference. We note that Spray is unrelated to Surrogate Variable Analysis [13, 14], a method developed to identify latent factors of variation present in microarray data.

For estimation, we implement a computationally efficient Expectation Conditional Maximization Either (ECME) algorithm [15, 16], which is adapted to fitting the association model in the presence of bilateral outcome missingness. The algorithm iterates between conditional maximization of the observed data log likelihood with respect to the regression parameters and conditional maximization of the EM objective function with respect to the covariance parameters. In addition, we derive the covariance estimators of all model parameters and implement a flexible Wald test for evaluating hypotheses about the target regression parameters.

We show analytically that the asymptotic relative efficiency of jointly modeling the target and surrogate outcomes, compared with marginally modeling the target outcome only, increases with the target missingness and the square of the target-surrogate correlation. We numerically demonstrate the analytical results through extensive simulations evaluating the empirical efficiency of the Spray Wald test.

Compared to complete case analysis, maximum likelihood estimation as implemented by Spray is efficient, making full use of the available data, and provides more precise estimates of the target regression parameters. All estimation and inference procedures described in this article have been implemented in an easy-to-use R package (`SurrogateRegression`), which is available on CRAN [17].

Using data from GTEx, we applied Spray to identify eQTL in the SSN, considering expression in blood, skeletal muscle, and the cerebellum as candidate surrogate outcomes. Compared with marginal eQTL mapping using expression in SSN only, Spray identified 4 to 5 times as many Bonferroni-significant eQTL, including all those identified by marginal analysis. Importantly, while the effect sizes estimated by Spray were nearly identical to those obtained via traditional, marginal eQTL mapping (*R*^2^ ≥ 0.995), the sampling variance of the estimates was reduced by up to 26%, on average, indicating that Spray increased power primarily by drawing on the correlated surrogate outcome to improve precision. Moreover, the effect sizes estimated by Spray are robust to the choice of surrogate outcome.

The remainder of this paper is organized as follows: Section 2 introduces the setting and model. Sections 3 and 4 detail the estimation and inference procedures. Section 5 addresses the estimand of Spray, and the asymptotic relative efficiency of jointly versus marginally modeling the target outcome. Section 6 presents the results of simulation studies, and Section 7 the application to the GTEx data. We conclude with discussions in Section 8.

## 2 Model and Setting

For each of *i* = 1, …, *n* independent subjects, suppose that two continuous outcomes are potentially observed: the *target outcome T_i_* and the *surrogate outcome S_i_*. Consider the model:

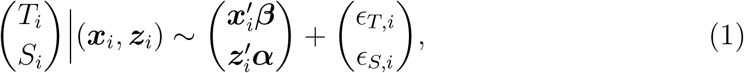

where ***x****_i_* is a *p* × 1 vector of covariates for the target outcome, with regression coefficients ***β***; ***z****_i_* is a *q* × 1 vector of covariates for the surrogate outcome, with regression coefficients ***α***; and ***ϵ***_i_ = (*ϵ_T,i_*, *ϵ_S,i_*)′ ~ *N*(**0, Σ**) with 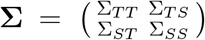. Let 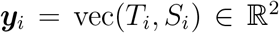 denote the 2 × 1 outcome vector, 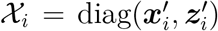 the 2 × (*p* + *q*) subject-specific design matrix, and ***γ*** = vec(***β***, ***α***) the (*p* + *q*) × 1 overall regression coefficient. With this notation, model (1) is succinctly expressible as: 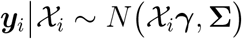.

Our derivations proceed under the assumption of residual normality. However, because in many applications, including eQTL mapping, the target and surrogate outcomes may be non-normal, we apply the rank-based inverse normal transformation (INT) to each outcome prior to analysis [18]. Application of INT, which ensures that the marginal distribution of each outcome is univariate normal, is common in eQTL studies, including all published analyses from GTEx [8]. While marginal normality of each outcome does not guarantee bivariate normality, our simulation studies demonstrate that this strategy provides unbiased estimation and valid inference even under residual distributions that are far from bivariate normal.

Unbiased estimation of model parameters requires that the target and surrogate outcomes are missing as random (MAR). For the *i*th subject, define the target *R_T,i_* and surrogate *R_S,i_* responses indicators:

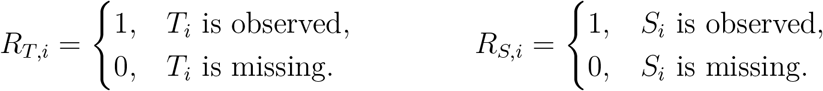

These indicators partition the *n* subjects into 3 missingness patterns: *complete cases* (*R_T,i_* = 1 and *R_S,i_* = 1); subjects with *target missingness* (*R_T,i_* = 0 and *R_S,i_* = 1); and subjects with *surrogate missingness* (*R_T,i_* = 1 and *R_S,i_* = 0). Subjects with neither outcome observed (*R_T,i_* = 0 and *R_S,i_* = 0) make no likelihood contribution and are not considered further. Supposing *n*_0_ complete cases, *n*_1_ subjects with target missingness, and *n*_2_ subjects with surrogate missingness, the total sample size is *n* = *n*_0_ + *n*_1_ + *n*_2_.

MAR requires that observation of the target outcome (*R_T,i_*) is unrelated to its value (*T_i_*), given the remaining data (*S_i_, **x**_i_, **z**_i_*), and likewise that *R_S,i_* is supposed unrelated to *S_i_*, given (*T_i_, **x**_i_, **z**_i_*). In our analysis of GTEx, the MAR assumption is plausible because donors were selected to be free of major diseases and the collection of tissue specimen was based on factors such as provision of consent and on the availability of sufficient tissue from the autopsy or surgical procedure [19, 20]. Importantly, the decision to ascertain a tissue sample was not directly based on gene expression.

## 3 Estimation

### 3.1 Regression Parameters

Define the response indicator matrix *R_i_* = diag(*R_T,i_, R_S,i_*), and note that *R_i_* is a projection matrix. The distribution of the observed data is expressible as:

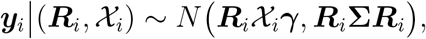

and the observed data log likelihood is:

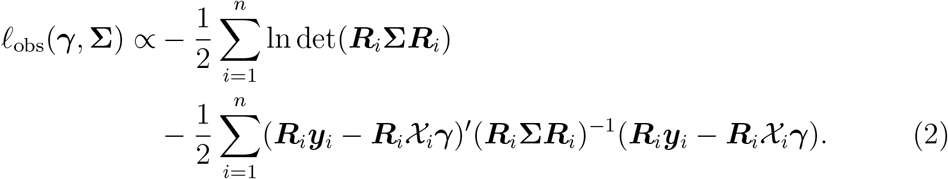

The observed data score equation for the regression parameters ***γ*** is:

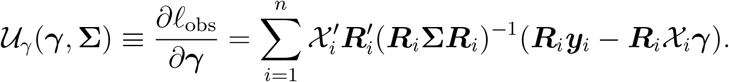

Conditional on **Σ**, the maximum likelihood estimator (MLE) of ***γ*** is the generalized least squares (GLS) estimator:

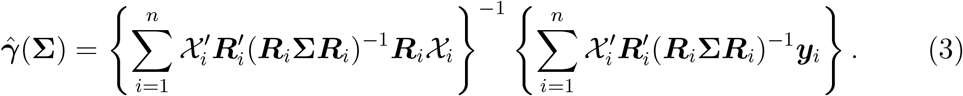

### 3.2 Covariance Matrix

Let *ϵ_i_* = (*y_i_* − *χ_iγ_*) denote the residual vector. The observed data score equation for **Σ** is:

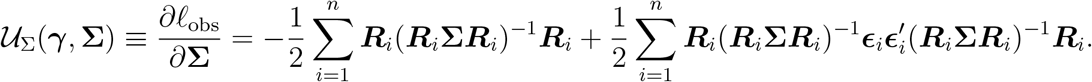

However, the score equation for **Σ** does not admit a closed form. To obtain the MLE, we apply the ECME algorithm [15, 16]. Define the 2 × 2 *residual outer product* matrix:

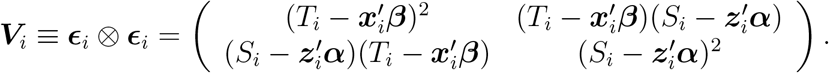

The *complete data log likelihood* is now expressible as:

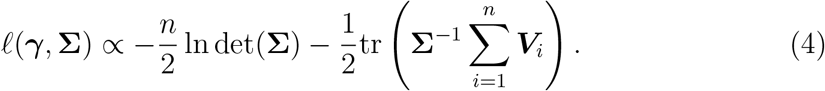

The *EM objective* is the expectation of the complete data log likelihood in (4) given the observed data 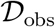 and the current parameter estimates (***γ***^(*r*)^, **Σ**^(*r*)^):

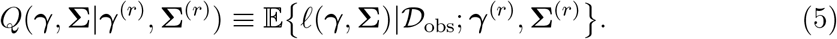

To obtain an expression for (5), define the *working outcome vector*:

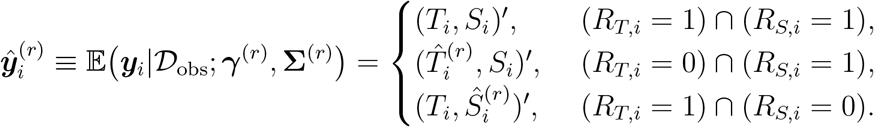

For complete cases, the working outcome vector is identically the observed outcome vector. For subjects with target missingness, the unobserved value of *T_i_* is replaced by its conditional expectation given the surrogate outcome and covariates:

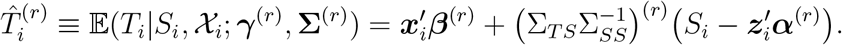

Note that we adopt the convention that 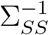 refers to subsetting the (*S, S*)th element of **Σ** then taking its inverse, as opposed to subsetting the (*S, S*)th of **Σ**^−1^. For subjects with surrogate missingness, the unobserved value of *S_i_* is replaced by its conditional expectation give the target outcome and covariates:

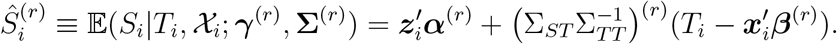

Let **Λ** = **Σ**^−1^ denote the precision matrix. Define the *working residual outer product*:

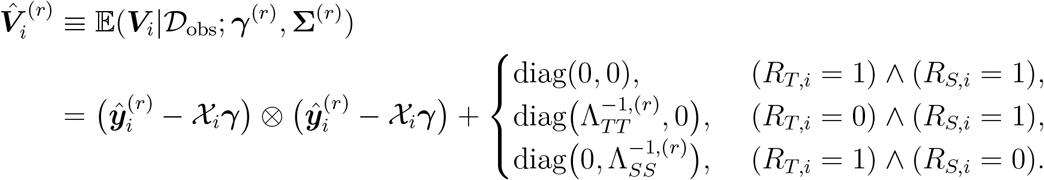

Expressed in terms of the working residual outer product, the EM objective function is:

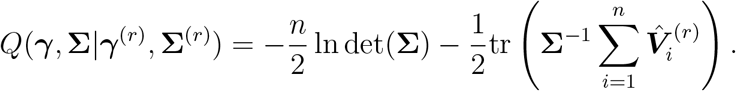

The EM score equation for **Σ** is:

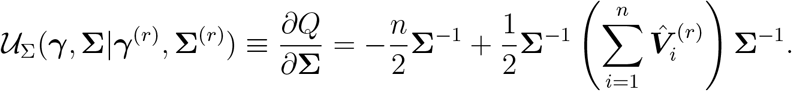

Conditional on ***γ***, the EM update for **Σ** is:

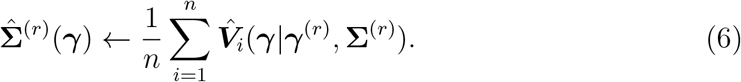

### 3.3 Optimization

Spray implements the following ECME algorithm, in which the regression parameters *γ* are updated via conditional maximization of the observed data log likelihood in (2), and the covariance matrix **Σ** is updated via conditional maximization of the EM objective in (5).

**Algorithm 1.**
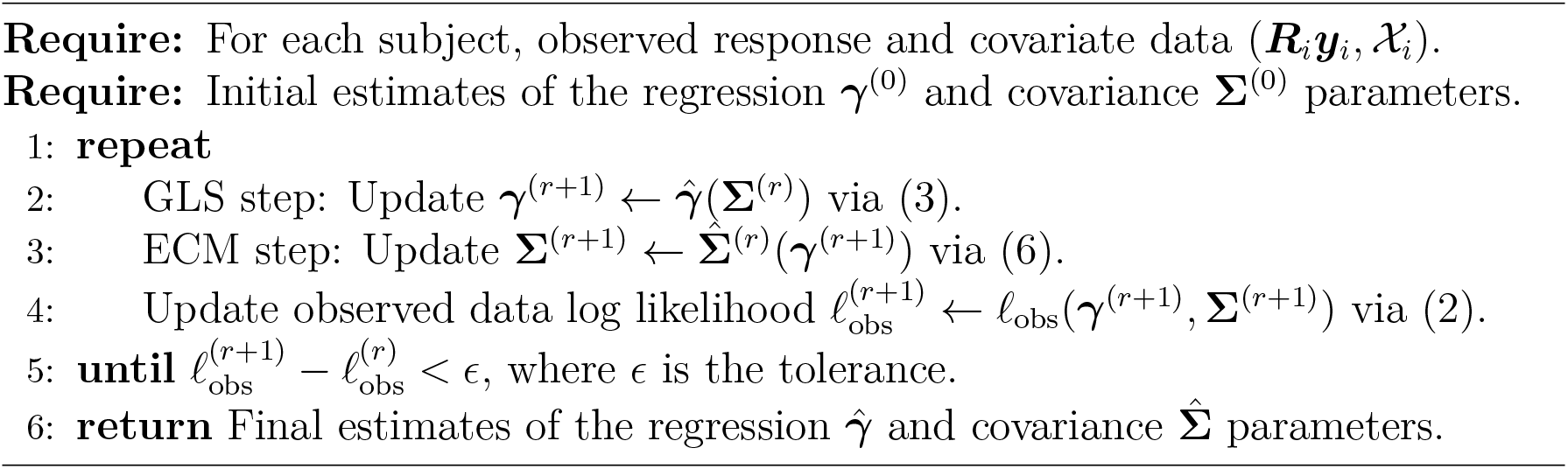
ECME for Bivariate Normal Regression

The accompanying `R` package initializes ***γ*** via ordinary least squares using all observed data:

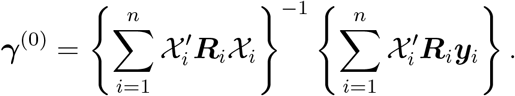

Given ***γ***^(0)^, **Σ** is initialized using the residual outer product of the *n*_0_ complete cases:

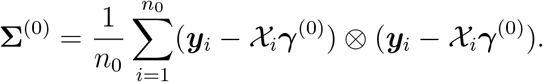

## 4 Inference

The ECME algorithm presented in the previous section does not provide the asymptotic information of the MLEs. The observed-data information matrices were obtained using the following identity:

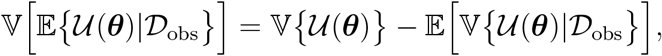

where 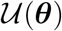 is the complete-data score, and 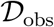 is the observed data. The observed-data information for the regression parameters ***γ*** decomposes as:

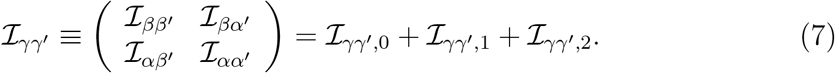

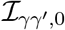 is the contribution of complete cases and takes the form:

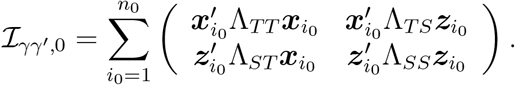

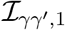 is the contribution of subjects with target missingness and 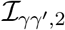 is the contribution of subjects with surrogate missingness; these take the following forms respectively:

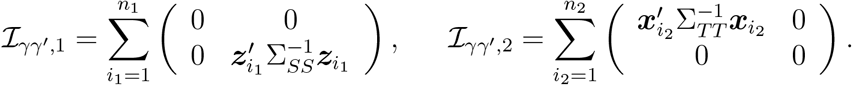

Complete cases contribute to the information for all regression parameters. Subjects with target missingness contribute to the information for the surrogate regression parameters ***α*** only, while subjects with surrogate missingness contribute to the information for the target regression parameters ***β*** only.

The observed-data information matrix for the covariance parameters (Σ*_TT_*, Σ*_TS_*, Σ*_SS_*) is presented in the supporting information, and follows a similar pattern of contributions. The cross information 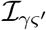 between the regression ***γ*** and covariance ***ς*** parameters is zero. Thus, the MLEs 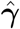 and 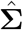 are asymptotically independent. For eQTL mapping, inference on the target regression parameter ***β* ⊆ *γ*** is performed using the standard Wald test, the details of which are also presented in the supporting information. Standard errors for all model parameters are provided by the accompanying R package, allowing for inference on ***α*** and **Σ** in addition to ***β***.

## 5 Analytical Considerations

### 5.1 Marginal Interpretation of the Regression Parameter

The choice to jointly model the target and surrogate outcomes, rather than conditioning on the surrogate to predict the target, has important ramifications when interpreting the regression parameters estimated by Spray. For exposition, suppose (1) is the generative model, and consider the setting where the target and surrogate means each depend on genotype *g_i_* only:

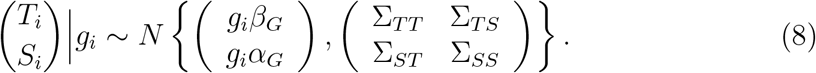

The implied marginal distribution of the target outcome is:

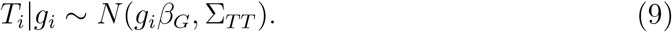

Observe that the regression parameter for genotype (*β_G_*) from the joint model (8) is identical to that appearing in the marginal model (9). This equality is unchanged by the presence or absence of an association *α_G_* between genotype *g_i_* the surrogate outcome *S_i_*. Importantly, as is confirmed by our simulation studies, this implies that inference on *β_G_* under the joint model (1) does not depend on the value of *α_G_*. The same is not true of a model that conditions on the surrogate outcome. In particular, when conditioning on the surrogate outcome, the target outcome is distributed as:

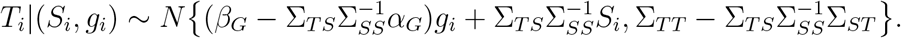

Suppose that the target and surrogate outcomes are associated (Σ*_TS_* ≠ 0), which is a prerequisite for modeling the surrogate outcome to improve inference on *β_G_*. Then, in a model that regresses *T_i_* on both (*S_i_, g_i_*), the magnitude and direction of the regression coefficient for genotype 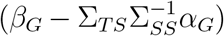 depends on whether and to what extent genotype is associated with the surrogate outcome (i.e. *α_G_*).

### 5.2 Efficiency Analysis

Consider again the genotype only model in (8). Suppose initially that all subjects are complete cases, and that the genotypes have been scaled such that: 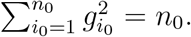. Under these assumptions, the *efficient information* for *β_G_* from (8) is:

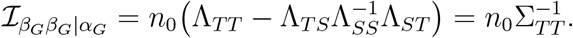

This is identical to the information for *β_G_* from the marginal model in (9). Thus, in the absence of missingness, inference on *β_G_* under the joint model (8) is asymptotically equivalent to inference on *β_G_* under the marginal model (9).

Now suppose there are *n*_0_ complete cases and *n*_1_ subjects with target missingness. For simplicity, assume no subjects have surrogate missingness, *n*_2_ = 0. Genotypes have again been scaled, within outcome missingness groups, such that 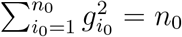 for complete cases and 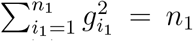 for subjects with target missingness. The efficient information from (8) becomes:

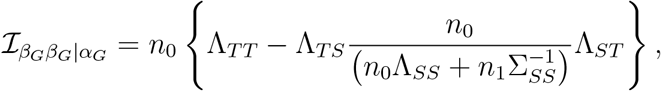

while the information for *β_G_* from the marginal model remains 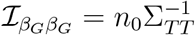. The *asymptotic relative efficiency* (ARE) of inference under the joint model (8) versus inference under marginal model (9) is:

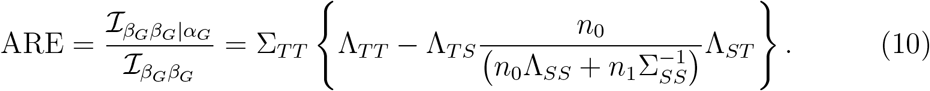

To better understand (10), suppose the covariance matrix in (8) is a correlation matrix, with Σ*_TT_* = Σ*_SS_* = 1, and correlation Σ*_TS_* = *ρ* ∈ (−1, 1). The ARE simplifies to:

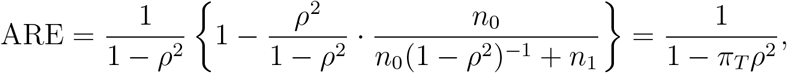

where *π_T_* = *n*_1_/(*n*_0_ + *n*_1_) is the proportion of subjects with target missingness. Now, if the target and surrogate outcomes are uncorrelated (*ρ* = 0), or if there is no target missingness (*π_T_* = 0), then the ARE is 1, and inference based on the marginal model is asymptotically equivalent to inference based on the joint model. For fixed target missingness *π_T_*, the ARE increases monotonically in the squared target-surrogate correlation *ρ*^2^. In the limit as *ρ* → 1, the ARE is maximized at (1 − *π_T_*)^−1^ = 1 + *n*_1_/*n*_0_. Likewise, for fixed target-surrogate correlation *ρ*^2^, the ARE increases monotonically in the target missingness *π_T_*. In the limit as *π_T_* → 1, which occurs when *n*_1_, the ARE is maximized at (1 − *ρ*^2^)^−1^. Overall, the power gain attributable to jointly modeling the target and surrogate outcomes is expected to increase with the squared target-surrogate correlation *ρ*^2^, and with the number of subjects with target missingness *n*_1_. This demonstrates an interesting property of the surrogate model: by leveraging the target-surrogate correlation, inference on the target outcome can be improved by incorporating information from subjects whose target outcomes are missing.

## 6 Simulation Studies

### 6.1 Brief Methods

The simulation methods are described in detail in the supporting information. Briefly, for each subject, the target *T_i_* and surrogate *S_i_* outcomes were simulated to depend on genotype *g_i_* and covariates *x_i_*, including age, sex, and genetic PCs. The simulations considered both normally and non-normally distributed residuals (*ϵ_T,i_*, *ϵ_S,i_*). INT was always applied to *T_i_* and *S_i_* prior to analysis. The number of complete cases was fixed at *n*_0_ = 10^3^. The numbers of subjects with missing outcomes (*n*_1_, *n*_2_) were varied to change the proportions (*π_T_, π_S_*) of subjects with target and surrogate missingness. Seven (target, surrogate) missingness patterns (*π_T_*, *π_S_*) were considered: no missingness (0.00, 0.00); unilateral target missingness {(0.25, 0.00), (0.50, 0.00), (0.75, 0.00)}; and bilateral outcome missingness {(0.25, 0.25), (0.50, 0.25), (0.25, 0.50)}. For each missingness pattern, the target-surrogate correlation *ρ* spanned {0.00, 0.25, 0.50, 0.75}.

### 6.2 Estimation

Table 1 considers estimation of the target genetic effect *β_G_*, the target variance Σ*_TT_*, and the target-surrogate correlation *ρ* both in the absence of missingness and in the presence of unilateral missingness in the target outcome. In all cases, parameter estimation was essentially unbiased, and the model-based standard errors (SEs), obtained from equation (7), agreed closely with the empirical standard deviations of the point estimates. Analogous tables for estimation of (*β_G_*, Σ*_TT_, ρ*) in the presence of bilateral missingness (S1), and for estimation of (*α_G_*, Σ*_SS_*) in the presence of both unilateral and bilateral missingness (S2) are presented in the supporting information.

**Table 1:**
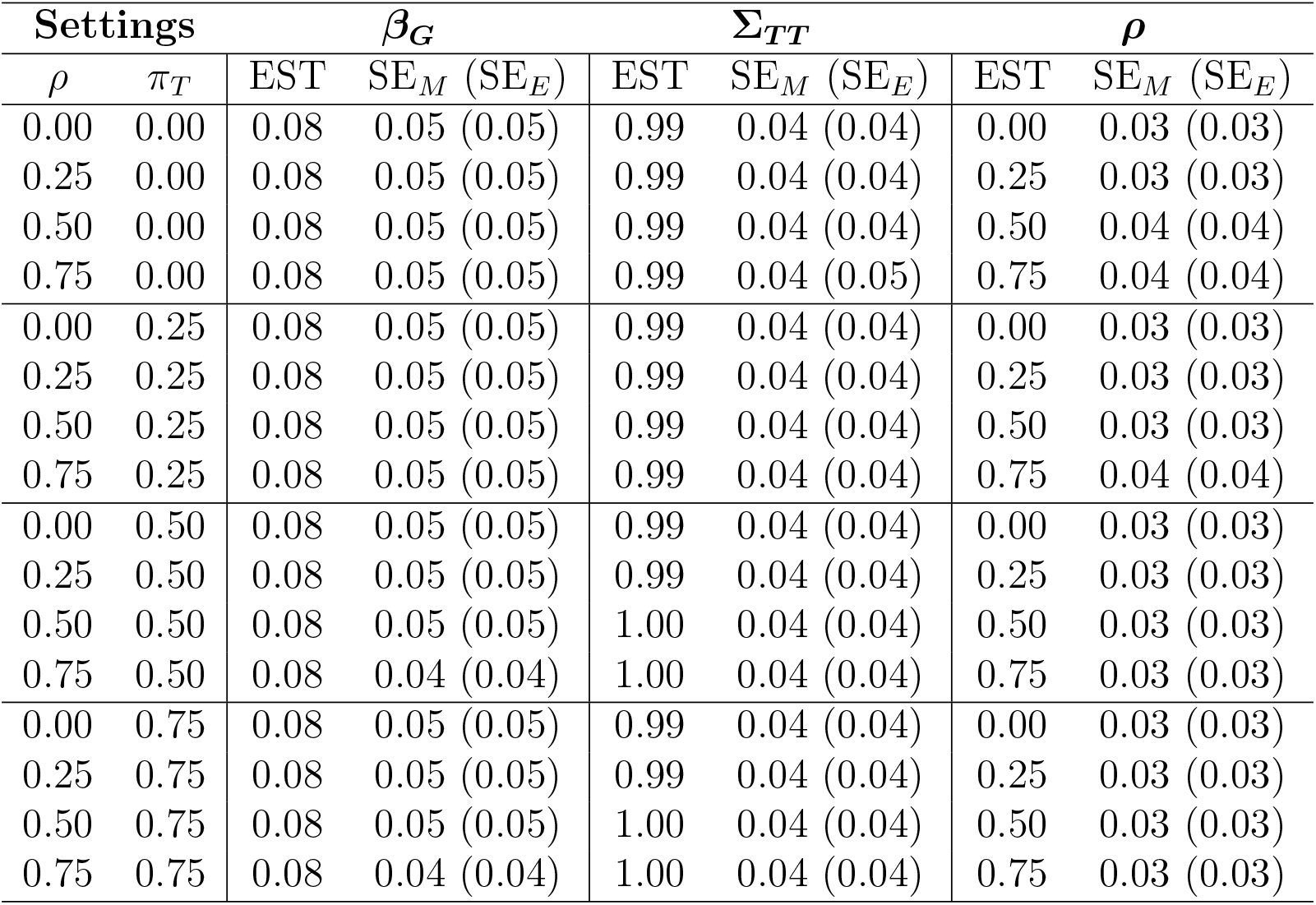
Target parameter estimation and standard error calibration across *R* = 5 × 10^7^ simulations in the presence of unilateral missingness. The number of complete cases was *n*_0_ = 10^3^. The true regression coefficient (*β_G_* ≈ 0.08) was chosen such that the heritability of the target outcome was 0.5% and the true variance of the target outcome was Σ*_TT_* = 1.00. The surrogate missingness was fixed at *π_S_* = 0.00 while the target missingness *π_T_* and target-surrogate correlation *ρ* were varied. The point estimate (EST) is the average across simulation replicates. The standard error is presented as the root mean square model-based standard error (SE*_M_*), followed by the empirical standard error (SE_*E*_) in parentheses, which is the standard deviation of the simulation point estimates.

| Settings | | $\beta_G$ | | $\Sigma_{TT}$ | | $\rho$ | |
| --- | --- | --- | --- | --- | --- | --- | --- |
| $\rho$ | $\pi_T$ | EST | $SE_M$ ($SE_E$) | EST | $SE_M$ ($SE_E$) | EST | $SE_M$ ($SE_E$) |
| 0.00 | 0.00 | 0.08 | 0.05 (0.05) | 0.99 | 0.04 (0.04) | 0.00 | 0.03 (0.03) |
| 0.25 | 0.00 | 0.08 | 0.05 (0.05) | 0.99 | 0.04 (0.04) | 0.25 | 0.03 (0.03) |
| 0.50 | 0.00 | 0.08 | 0.05 (0.05) | 0.99 | 0.04 (0.04) | 0.50 | 0.04 (0.04) |
| 0.75 | 0.00 | 0.08 | 0.05 (0.05) | 0.99 | 0.04 (0.05) | 0.75 | 0.04 (0.04) |
| 0.00 | 0.25 | 0.08 | 0.05 (0.05) | 0.99 | 0.04 (0.04) | 0.00 | 0.03 (0.03) |
| 0.25 | 0.25 | 0.08 | 0.05 (0.05) | 0.99 | 0.04 (0.04) | 0.25 | 0.03 (0.03) |
| 0.50 | 0.25 | 0.08 | 0.05 (0.05) | 0.99 | 0.04 (0.04) | 0.50 | 0.03 (0.03) |
| 0.75 | 0.25 | 0.08 | 0.05 (0.05) | 0.99 | 0.04 (0.04) | 0.75 | 0.04 (0.04) |
| 0.00 | 0.50 | 0.08 | 0.05 (0.05) | 0.99 | 0.04 (0.04) | 0.00 | 0.03 (0.03) |
| 0.25 | 0.50 | 0.08 | 0.05 (0.05) | 0.99 | 0.04 (0.04) | 0.25 | 0.03 (0.03) |
| 0.50 | 0.50 | 0.08 | 0.05 (0.05) | 1.00 | 0.04 (0.04) | 0.50 | 0.03 (0.03) |
| 0.75 | 0.50 | 0.08 | 0.04 (0.04) | 1.00 | 0.04 (0.04) | 0.75 | 0.03 (0.03) |
| 0.00 | 0.75 | 0.08 | 0.05 (0.05) | 0.99 | 0.04 (0.04) | 0.00 | 0.03 (0.03) |
| 0.25 | 0.75 | 0.08 | 0.05 (0.05) | 0.99 | 0.04 (0.04) | 0.25 | 0.03 (0.03) |
| 0.50 | 0.75 | 0.08 | 0.05 (0.05) | 1.00 | 0.04 (0.04) | 0.50 | 0.03 (0.03) |
| 0.75 | 0.75 | 0.08 | 0.04 (0.04) | 1.00 | 0.04 (0.04) | 0.75 | 0.03 (0.03) |

To evaluate sensitivity of the estimation procedure to the bivariate normality assumption, additional simulations were conducted in which the target and surrogate residuals were generated from non-normal distributions, including bivariate versions of the exponential, log-normal, and Student *t*_3_ distributions. The bias and SE for estimating the parameter of primary interest, target genetic effect *β_G_*, are presented in supporting table (S3). Even when applied to skewed and kurtotic phenotypes, the estimation procedure remained unbiased and the SEs correctly calibrated, suggesting robustness to the residual distribution.

### 6.3 Type I Error Simulations

Table 2 presents the empirical type I error and non-centrality parameter (NCP) of the Spray Wald test in the presence of unilateral missingness; estimates under bilateral missingness are presented in supporting table S4. For these simulations the genetic effects were set to zero (*β_G_* = 0.00) and the null hypothesis *H*_0_ : *β_G_* = 0 was evaluated. The type I error was controlled to within 0.8% of the nominal level, and the NCP was within 0.2% of the reference value; both were insensitive to outcome missingness and target-surrogate correlation. Supporting figures S1-S2 demonstrate that, across outcome missingness patterns and target-surrogate correlation levels, the p-values provided by the Spray Wald test were uniformly distributed under the null. Thus, Spray provides a valid test of association between genotype and the target outcome. Supporting tables S5-S7 and figures S3-S5 indicate that the type I error is well-controlled even when the distribution of the phenotypic residuals is non-normal. Supporting table S8 verifies that control of the type I error becomes increasingly tight as sample size increases, to within 0.2% of nominal by a sample size of 20 × 10^3^.

**Table 2:**
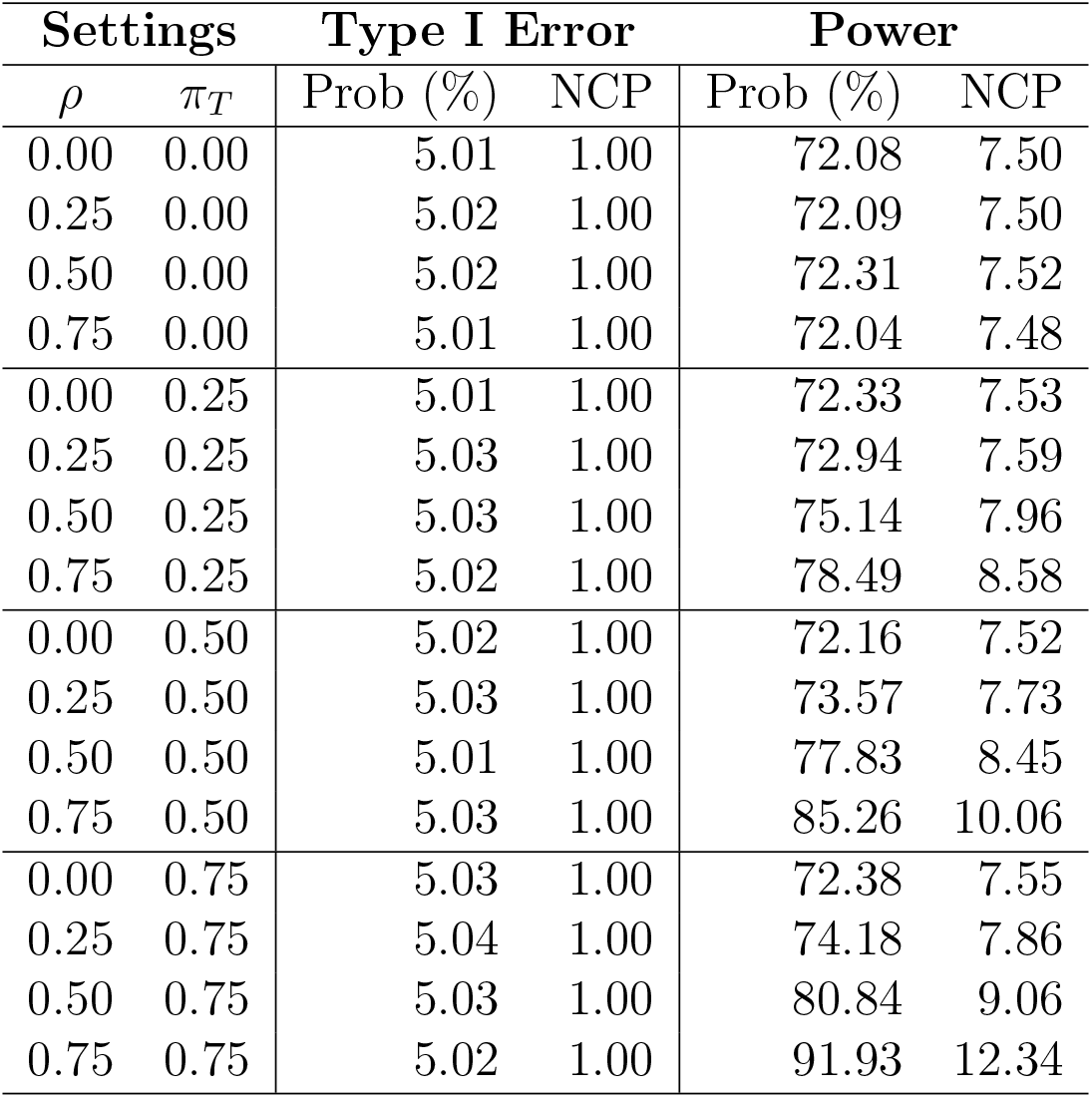
Empirical type I error and power of the Spray Wald test across *R* = 5 × 10^7^ simulation replicates in the presence of unilateral missingness. The number of complete cases was *n*_0_ = 10^3^. The surrogate missingness was fixed at *π_S_* = 0. For type I error, *β_G_* = 0 while for power *β_G_* was selected to explain 0.5% of variation in the target outcome. The target missingness *π_T_* and target-surrogate correlation *ρ* were varied. Prob refers to the rejection probability at a target type I error of 5% and NCP is the non-centrality parameter of the Wald test.

| Settings |  | Type I Error |  | Power |  |
| --- | --- | --- | --- | --- | --- |
| $\rho$ | $\pi_T$ | Prob (%) | NCP | Prob (%) | NCP |
| 0.00 | 0.00 | 5.01 | 1.00 | 72.08 | 7.50 |
| 0.25 | 0.00 | 5.02 | 1.00 | 72.09 | 7.50 |
| 0.50 | 0.00 | 5.02 | 1.00 | 72.31 | 7.52 |
| 0.75 | 0.00 | 5.01 | 1.00 | 72.04 | 7.48 |
| 0.00 | 0.25 | 5.01 | 1.00 | 72.33 | 7.53 |
| 0.25 | 0.25 | 5.03 | 1.00 | 72.94 | 7.59 |
| 0.50 | 0.25 | 5.03 | 1.00 | 75.14 | 7.96 |
| 0.75 | 0.25 | 5.02 | 1.00 | 78.49 | 8.58 |
| 0.00 | 0.50 | 5.02 | 1.00 | 72.16 | 7.52 |
| 0.25 | 0.50 | 5.03 | 1.00 | 73.57 | 7.73 |
| 0.50 | 0.50 | 5.01 | 1.00 | 77.83 | 8.45 |
| 0.75 | 0.50 | 5.03 | 1.00 | 85.26 | 10.06 |
| 0.00 | 0.75 | 5.03 | 1.00 | 72.38 | 7.55 |
| 0.25 | 0.75 | 5.04 | 1.00 | 74.18 | 7.86 |
| 0.50 | 0.75 | 5.03 | 1.00 | 80.84 | 9.06 |
| 0.75 | 0.75 | 5.02 | 1.00 | 91.93 | 12.34 |

It is important to note that throughout the simulations, the target-surrogate correlation was estimated. For a given realization of the data, the MLE 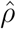 will differ from 0 even when in truth *ρ* = 0. The type I error simulations verify that this spurious estimated correlation does not compromise inference on *β_G_*.

### 6.4 Power Simulations

Table 2 presents the estimated power and NCP of the `Spray` Wald test for rejecting the *H*_0_ : *β_G_* = 0 in the presence of unilateral missingness; estimates under bilateral missingness are presented in supporting table S4. For these simulations, *β_G_* was chosen such that the proportion of variation in the target outcome explained by variation in genotype (i.e. the heritability) was 0.5%. Figures 1 and S6 present power curves describing how the probability of correctly rejecting the null hypothesis increases as the heritability increases from 0.1% to 1.0%. In the absence of target missingness, no additional power was gained by modeling the surrogate outcome. In the presence of target missingness, the power of the Spray Wald test increased with the target-surrogate correlation, and the relative improvement increased with the extent of target missingness. Supporting tables S5-S7 and figures S7-S9 demonstrate that similar trends with respect to power held under model misspecification. Whereas power under an exponential data generating process nearly matched that under a normal data generating process, power was attenuated in the more kurtotic cases of log-normal and Student *t*_3_ residuals.

**Figure 1:**
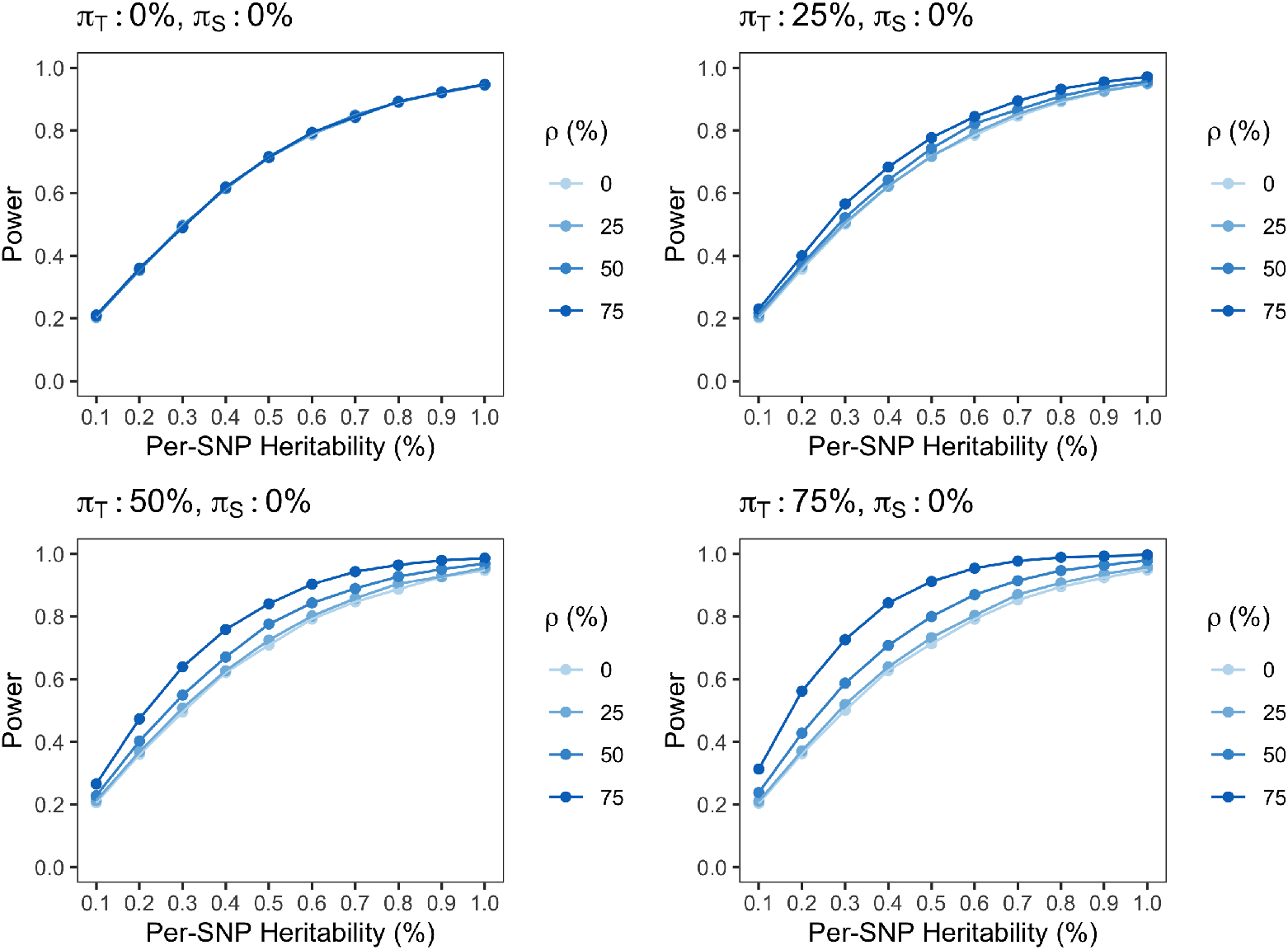
Power curves for the Spray test of association in the presence of unilateral missingness. The number of complete cases was *n*_0_ = 10^3^, and the type I error was *α* = 0.05. Each point on the curve is the average across *R* = 5 × 10^5^ simulation replicates. The standard errors of the point estimates were negligible. The target regression coefficient *β_G_* was varied between 0.037 and 0.14 to achieve heritabilities between 0.1% and 1.0%, while the surrogate regression coefficient *α_G_* was fixed at zero. The surrogate missingness was held at *π_S_* = 0, while the target missingness *π_T_* and target-surrogate correlation *ρ* were varied. Note that this figure appears in color in the electronic version of this article, and any mention of color refers to that version.

### 6.5 Empirical Relative Efficiency

To validate the ARE formula in equation (10), we conducted simulations comparing the Spray estimator 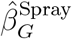 of *β_G_* with the marginal estimator 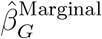 from the model:

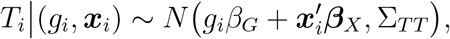

These simulations quantify the efficiency gain attributable to incorporating information from the surrogate. Table 3 compares the empirical variances of 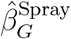 and 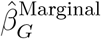 in the presence of unilateral missingness, while supporting table S9 compares the empirical variances under bilateral missingness. In the absence of target missingness (*π_T_* = 0), or when the target-surrogate correlation was zero (*ρ* = 0), the empirical RE was one, as predicted by (10). Thus, while jointly modeling the target and surrogate outcomes is unnecessary in the absence of missingness, power is not substantially diminished by modeling an uninformative surrogate. In the presence of missingness, modeling an uninformative surrogate (*ρ* = 0) did not spuriously inflate the RE. As the target missingness (*π_T_*) and target-surrogate correlation (*ρ*) increased, the empirical RE increased as predicted by (10). The precise agreement between the empirical and theoretical REs suggests that equation (10) could prove useful for study design.

**Table 3:**
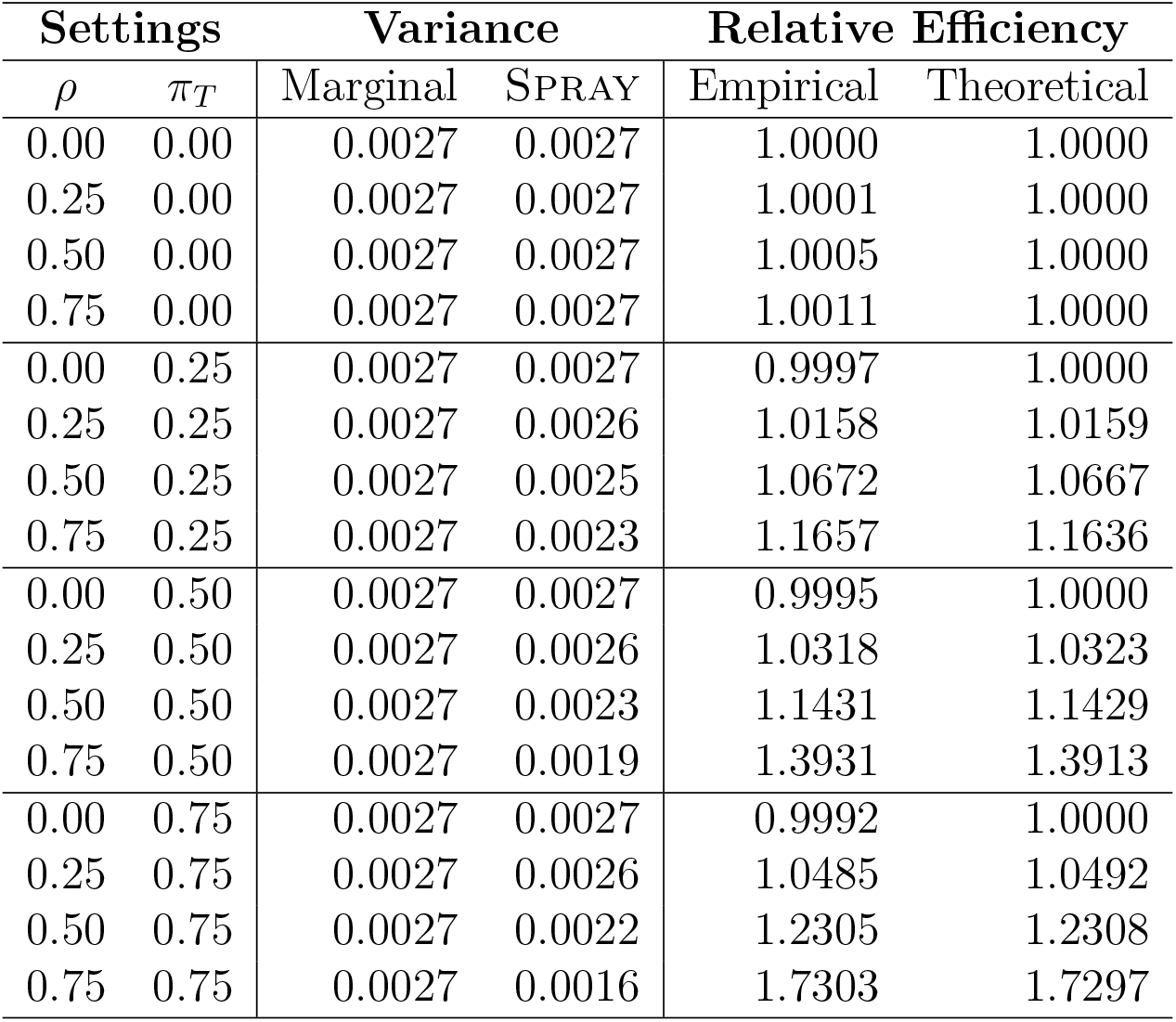
Empirical relative efficiency comparing the Spray estimator to the marginal estimator of *β_G_* test across *R* = 5 × 10^7^ simulation replicates in the presence of unilateral missingness. The number of complete cases was *n*_0_ = 10^3^. The true regression coefficient (*β_G_* ≈ 0.08) was chosen such that the heritability of the target outcome was 0.5%. The true variances of the target and surrogate outcomes were Σ*_TT_* = Σ*_SS_* = 1.00. The surrogate missingness was fixed at *π_S_* = 0.00. The target missingness *π_T_* and target-surrogate correlation *ρ* were varied. Variance refers to the empirical variance of the corresponding estimator across simulation replicates. The empirical RE is the ratio of the variance of 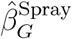 to that of 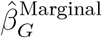. The theoretical RE was obtained from (10).

| Settings |  | Variance |  | Relative Efficiency |  |
| --- | --- | --- | --- | --- | --- |
| $\rho$ | $\pi_T$ | Marginal | SPRAY | Empirical | Theoretical |
| 0.00 | 0.00 | 0.0027 | 0.0027 | 1.0000 | 1.0000 |
| 0.25 | 0.00 | 0.0027 | 0.0027 | 1.0001 | 1.0000 |
| 0.50 | 0.00 | 0.0027 | 0.0027 | 1.0005 | 1.0000 |
| 0.75 | 0.00 | 0.0027 | 0.0027 | 1.0011 | 1.0000 |
| 0.00 | 0.25 | 0.0027 | 0.0027 | 0.9997 | 1.0000 |
| 0.25 | 0.25 | 0.0027 | 0.0026 | 1.0158 | 1.0159 |
| 0.50 | 0.25 | 0.0027 | 0.0025 | 1.0672 | 1.0667 |
| 0.75 | 0.25 | 0.0027 | 0.0023 | 1.1657 | 1.1636 |
| 0.00 | 0.50 | 0.0027 | 0.0027 | 0.9995 | 1.0000 |
| 0.25 | 0.50 | 0.0027 | 0.0026 | 1.0318 | 1.0323 |
| 0.50 | 0.50 | 0.0027 | 0.0023 | 1.1431 | 1.1429 |
| 0.75 | 0.50 | 0.0027 | 0.0019 | 1.3931 | 1.3913 |
| 0.00 | 0.75 | 0.0027 | 0.0027 | 0.9992 | 1.0000 |
| 0.25 | 0.75 | 0.0027 | 0.0026 | 1.0485 | 1.0492 |
| 0.50 | 0.75 | 0.0027 | 0.0022 | 1.2305 | 1.2308 |
| 0.75 | 0.75 | 0.0027 | 0.0016 | 1.7303 | 1.7297 |

## 7 Application to Identifying SSN eQTL in GTEx

### 7.1 Brief Data Analysis Methods

Details of the GTEx analysis are presented in the supporting information. Briefly, gene expression in SSN was the target outcome. Three surrogate analyses were conduct in parallel, based respectively on whole blood, skeletal muscle, and cerebellum as the surrogate. We address the idea of using multiple surrogates simultaneously in the Discussion. For inclusion in the analysis, a transcript was required to be expressed in both the target and surrogate tissues. SNPs in *cis* to an expressed transcript were tested for association. Two associations methods were applied, a marginal analysis that regresses the target outcome only on genotype and covariates, and a joint analysis (Spray) that regresses the target and surrogate outcomes on genotype and covariates. Significance was declared at the Bonferroni threshold, adjusted for the number of SNP-transcript pairs tested for association.

### 7.2 Results

There were 80 genotyped subjects with expression in SSN. Supporting table S10 presents the sample sizes available in the 3 candidate surrogate tissues. The total sample size was largest for muscle (*n* = 507) and smallest for cerebellum (*n* = 168). However, as figure S10 demonstrates, the correlation between cerebellum and SSN was typically higher than that between muscle or blood and SSN. The root-mean-square correlation between the target and surrogate tissues was 0.18 for blood and muscle in comparison to 0.31 for cerebellum. Moreover, the number of transcripts expressed in both SSN and the surrogate tissue was greatest for cerebellum (table S11).

Table 4 compares the marginal and joint (Spray) eQTL analyses of SSN by surrogate tissue. In all cases, joint analysis identified more Bonferroni significant associations and did so more efficiently. All eQTL identified by the marginal analysis were also identified by the joint analysis, but not conversely. Most eQTL were detected when using cerebellum as the surrogate, although muscle in fact provided a more efficient surrogate, meaning the estimated standard errors were on average lower. More eQTL were identified with cerebellum because 19 transcripts containing 33 significant eQTL were expressed in cerebellum but not muscle.

**Table 4:** Comparison of marginal and Spray analyses by surrogate tissue. Significant eQTL were identified at the Bonferroni threshold for each analysis. The mean *χ*^2^ statistic is calculate across those eQTL significant under either the marginal or joint (Spray) analyses. Relative efficiency is calculated as the mean of the ratio of the sampling variance of the marginal estimator to the Spray estimator.

| Surrogate | Significant eQTL | | Mean $\chi^2$ | | Relative Efficiency |
| --- | --- | --- | --- | --- | --- |
|  | Marginal | SPRAY | Marginal | SPRAY |  |
| Blood | 24 | 111 | 40.8 | 48.0 | 1.18 |
| Muscle | 37 | 149 | 40.8 | 50.1 | 1.26 |
| Cerebellum | 42 | 176 | 40.3 | 49.1 | 1.25 |

Figure 2A compares the estimated effect sizes of the marginal and joint analyses using cerebellum as the surrogate. Analogous figures for blood and muscle are presented in supporting figures S11 and S12. In all cases, the effect sizes were tightly correlated, verifying that Spray estimates the same effect as traditional, marginal analyses. However, figure 2B demonstrates that Spray provides greater power to detect eQTL. From table 4, this is because, at eQTL considered significant by either marginal or joint analysis, Spray provided standard errors that were up to 26% smaller, on average. Finally, figure 3 considers the concordance in effect sizes and p-values among SNP-transcript pairs that were tested for association in at least 2 of the surrogate analyses and were significant in at least 1. The tight correlation in effect sizes suggests that Spray is robust to the choice of surrogate outcome.

**Figure 2:**
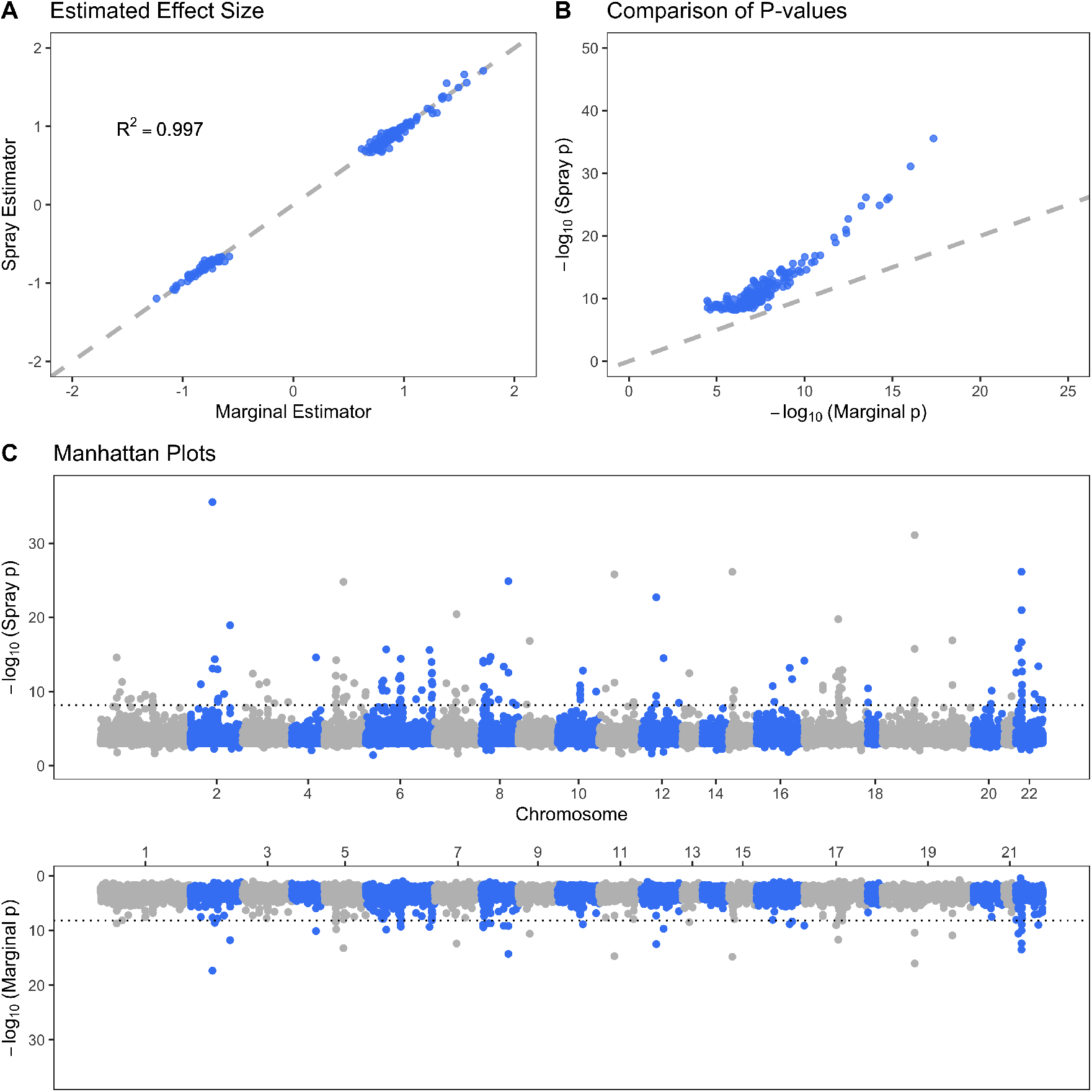
Comparison of the marginal and joint (Spray) eQTL analysies of substantia nigra, using cerebellum as the surrogate tissue. A. Estimated effect size from the joint analysis vs. the estimated effect size from the marginal analysis for eQTL significant in at least 1 of the analyses. B. P-value from the joint analysis vs. p-value from the marginal analysis for eQTL significant in at least 1 of the analyses. C. Mirrored Manhattan plots comparing the p-values of the joint and marginal analyses by genomic position. Dotted line is the Bonferroni significance threshold. Note that this figure appears in color in the electronic version of this article, and any mention of color refers to that version.

**Figure 3:**
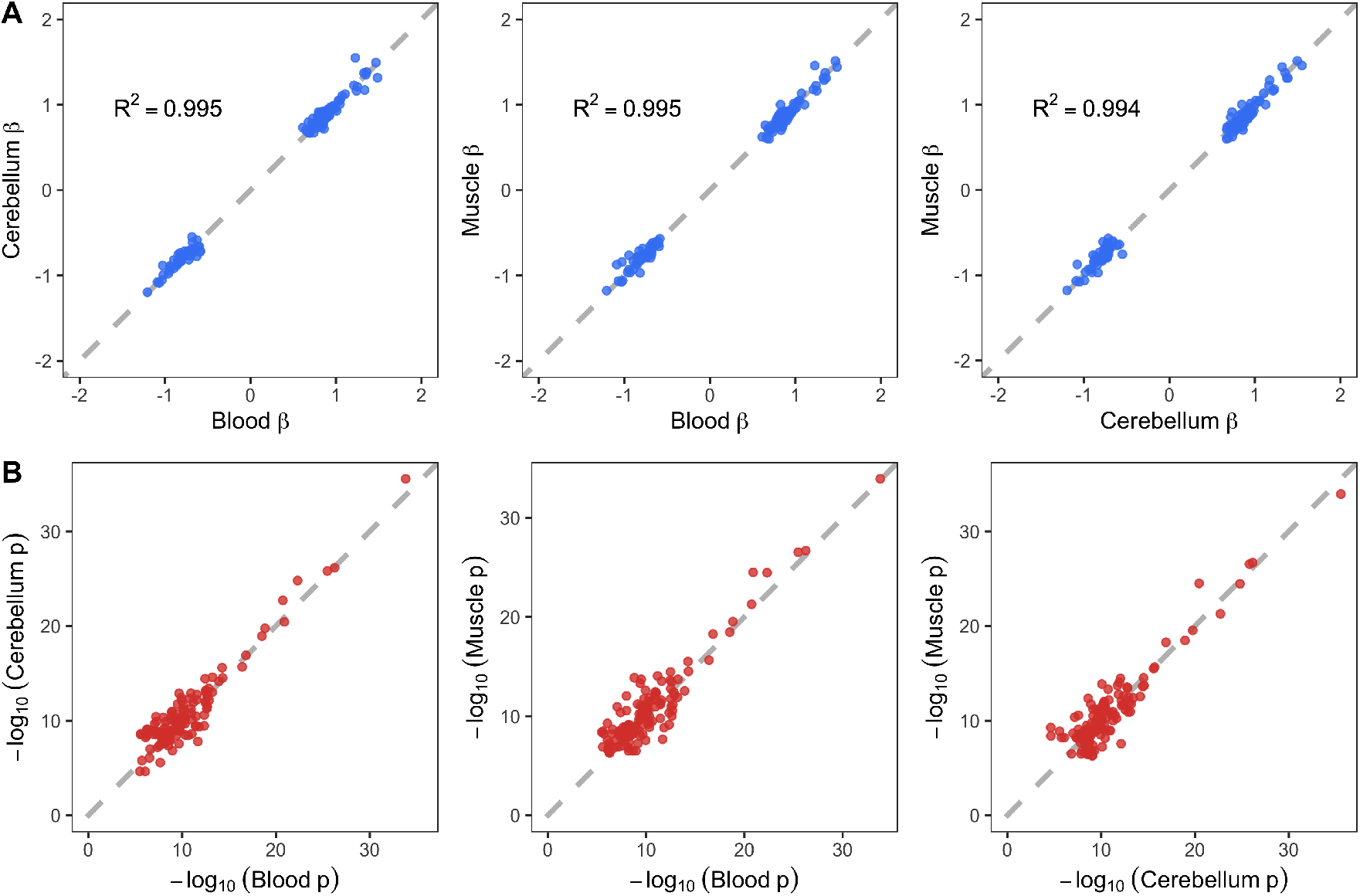
Effect of the surrogate outcome on the results of joint (Spray) analysis. A. Estimated effect size by surrogate outcome for eQTL evaluated in at least 2 of the surrogate analyses and significant in at least 1. B. Association p-value by surrogate outcome, again for QTL evaluated in at least 2 of the surrogate analyses and significant in at least 1. Note that this figure appears in color in the electronic version of this article, and any mention of color refers to that version.

## 8 Discussion

In this article, we have proposed leveraging a correlated surrogate outcome to improve inference on a partially missing target outcome, and derived a computationally efficient, ECME-type algorithm for fitting the association model. We demonstrated analytically and empirically, though extensive simulations and in real data, that the Spray test of association, which incorporates information from the target and surrogate outcomes, is more efficient than the marginal test of association, which incorporates information from the target outcome only. The efficiency of Spray increases with the target missingness, and with the square of the target-surrogate correlation. Moreover, we showed that modeling the surrogate as an outcome, rather than conditioning on it as a covariate, allows Spray to estimate the same effect size as traditional, marginal analysis. All estimation and inference procedures described in this article have been made available as an R package [17].

We applied Spray to eQTL mapping in GTEx, using expression in SSN as the target outcome and expression in one of blood, muscle, or cerebellum as the surrogate outcome. Relative to marginal analysis, joint analysis using Spray consistently identified more Bonferroni significant associations. Although the joint and marginal effect size estimates were highly concordant (*R*^2^ ≥ 0.995), the Spray estimator was up to 26.0% more efficient, on average, at Bonferroni-significant eQTL. The choice of surrogate tissue highlighted a trade-off between the quality of the surrogate, as measured by its correlation with the target outcome, and the availability of the surrogate. Expression in muscle was available for 3 times as many subjects as expression in cerebellum, yet expression in cerebellum was better correlated with expression in SSN. Although the effect size estimated by Spray is unaffected by the choice of surrogate, the power is; sample sizes being equal, the better correlated surrogate is preferred. When the available sample sizes are not equal, equation (10) may be used to examine the trade-off.

Our work suggests several areas for further improvement. Although INT was applied to ensure marginal normality of the target and surrogate outcomes, joint bivariate normality is not guaranteed. While our results show that INT confers robustness to residual non-normality, a future direction is to develop association tests that allow for arbitrary patterns of outcome missingness but do not require specification of a joint distribution. Instead of maximum likelihood based estimation, this procedure could use a set of inverse probability weighted estimating equations [21].

Another avenue for future development is to incorporate multiple surrogate outcomes. One way to achieve this would be to extend the bivariate normal regression framework to a multivariate normal regression framework. However, there are drawbacks to directly modeling multiple surrogate outcomes: the number of nuisance covariance parameters increases quadratically with the number of surrogates, and the number of potential missingness patterns increases exponentially. Finally, although the current work was motivated by eQTL mapping, the idea of leveraging a surrogate outcome to improve inference on a partially missing target outcome is broadly applicable. For example, in large cohort studies such as the UK Biobank [22], the target outcome may be any incompletely ascertained phenotype, such as the concentration of a biomarker only measured for a subset of participants, while the surrogate outcome may be a readily ascertained phenotype, such as a risk score based on diagnostic codes from electronic health records.

## Acknowledgments

This work was supported by the National Institutes of Health grants R35 CA197449 and F31 HL140822 (to Z.M.) and R35 CA197449, P01 CA134294, U01 HG009088, U01HG012064, 19 CA203654, and R01 HL113338 (to X.L.).

## Supporting Information

Additional simulation results are available in the Support Information published online. `Spray` is available at https://CRAN.R-project.org/package=SurrogateRegression. Code for reproducing all simulations and summary statistics from the GTEx analysis are available at: https://github.com/zrmacc/Surrogate-Replication-eQTL.

