## Supporting Information for "Leveraging a surrogate outcome to improve inference on a partially missing target outcome"

#### Contents

**1 Variance Component Information Matrix** **1**

**2 Wald Test of Association** **1**

**3 Detailed Simulations Methods** **2**

**4 Estimation** **3**

### 1 Variance Component Information Matrix

Let $\varsigma = \text{vec}(\Sigma_{TT}, \Sigma_{TS}, \Sigma_{SS})$ vectorize the covariance parameters. The observed-data information for $\boldsymbol{\varsigma}$ likewise decomposes as:

$$\mathcal{I}_{\varsigma\varsigma'} = \begin{pmatrix} \mathcal{I}_{\varsigma_1\varsigma_1} & \mathcal{I}_{\varsigma_1\varsigma_2} & \mathcal{I}_{\varsigma_1\varsigma_3} \\ \mathcal{I}_{\varsigma_2\varsigma_1} & \mathcal{I}_{\varsigma_2\varsigma_2} & \mathcal{I}_{\varsigma_2\varsigma_3} \\ \mathcal{I}_{\varsigma_3\varsigma_1} & \mathcal{I}_{\varsigma_3\varsigma_2} & \mathcal{I}_{\varsigma_3\varsigma_3} \end{pmatrix} = \mathcal{I}_{\varsigma\varsigma',0} + \mathcal{I}_{\varsigma\varsigma',1} + \mathcal{I}_{\varsigma\varsigma',2}. \tag{S1}$$

$\mathcal{I}_{\varsigma\varsigma',0}$ is the contribution of complete cases:

$$\mathcal{I}_{\varsigma\varsigma',0} = \frac{n_0}{2} \begin{pmatrix} \Lambda_{TT}^2 & 2\Lambda_{TT}\Lambda_{TS} & \Lambda_{TS}^2 \\ 2\Lambda_{TT}\Lambda_{TS} & 2\Lambda_{TS}^2 + 2\Lambda_{TT}\Lambda_{SS} & 2\Lambda_{TS}\Lambda_{SS} \\ \Lambda_{TS}^2 & 2\Lambda_{TS}\Lambda_{SS} & \Lambda_{SS}^2 \end{pmatrix}.$$

$\mathcal{I}_{\varsigma\varsigma',1}$ is the contribution of subjects with target missingness and $\mathcal{I}_{\varsigma\varsigma',1}$ is the contribution of subjects with surrogate missingness; they take the following forms respectively:

$$\mathcal{I}_{\varsigma\varsigma',1} = \frac{n_1}{2} \begin{pmatrix} 0 & 0 & 0 \\ 0 & 0 & 0 \\ 0 & 0 & \Sigma_{SS}^{-2} \end{pmatrix}, \quad \mathcal{I}_{\varsigma\varsigma',2} = \frac{n_2}{2} \begin{pmatrix} \Sigma_{TT}^{-2} & 0 & 0 \\ 0 & 0 & 0 \\ 0 & 0 & 0 \end{pmatrix}.$$

### 2 Wald Test of Association

Suppose that the target of inference is a subset $\boldsymbol{\beta}_A$ of regression parameters for the target outcome. The problem is symmetric if considering inference on a subset of parameters for the surrogate outcome. Partition $\boldsymbol{\beta} = (\boldsymbol{\beta}_A, \boldsymbol{\beta}_B)$, where $\boldsymbol{\beta}_A$ denotes the parameter of interest, and let $\boldsymbol{\eta} = (\boldsymbol{\beta}_B, \boldsymbol{\alpha})$ denote the remaining, nuisance regression parameters. The joint information of the target and nuisance regression parameters $(\boldsymbol{\beta}_A, \boldsymbol{\eta})$ is:

$$\mathcal{I}_{(\beta_A,\eta)(\beta_A,\eta)'} = \begin{pmatrix} \mathcal{I}_{\beta_A\beta_A'} & \mathcal{I}_{\beta_A\eta'} \\ \mathcal{I}_{\eta\beta_A'} & \mathcal{I}_{\eta\eta'} \end{pmatrix},$$

which is a permutation of the information matrix for $\boldsymbol{\gamma}$ in main text equation (7). The *efficient information* for $\boldsymbol{\beta}_A$ is: $\mathcal{I}_{\beta_A\beta_A|\eta} = \mathcal{I}_{\beta_A\beta_A'} - \mathcal{I}_{\beta_A\eta'}\mathcal{I}_{\eta\eta'}^{-1}\mathcal{I}_{\eta\beta_A'}$. Under the null hypothesis $H_0 : \boldsymbol{\beta}_A = \boldsymbol{\beta}_A^*$, the maximum likelihood estimate $\hat{\boldsymbol{\beta}}_A$ is asymptotically normal:

$$\left(\hat{\boldsymbol{\beta}}_A - \boldsymbol{\beta}_A^*\right) \dot{\sim} N\left(\mathbf{0}, \mathcal{I}_{\beta_A\beta_A'|\eta}^{-1}\right).$$

The Wald test of $H_0 : \boldsymbol{\beta}_A = \boldsymbol{\beta}_A^*$ is:

$$T_W = \left(\hat{\boldsymbol{\beta}}_A - \boldsymbol{\beta}_A^*\right)'\left(\mathcal{I}_{\beta_A\beta_A'|\eta}\right)\left(\hat{\boldsymbol{\beta}}_A - \boldsymbol{\beta}_A^*\right) \dot{\sim} \chi_\nu^2, \tag{S2}$$

For increasing $n$, $T_W$ follows a $\chi_\nu^2$ distribution with $\nu = \dim(\boldsymbol{\beta}_A)$ degrees of freedom. SPRAY uses the Wald statistic in (S2) for identification of eQTL in the target tissue.

### 3 Detailed Simulations Methods

Additively coded genotypes $g_i \in \{0, 1, 2\}$ were drawn from a binomial distribution with minor allele frequency of 25%. Covariates included age and sex: age was drawn from a gamma distribution with mean 50 and variance 10; sex was drawn from a Bernoulli distribution with proportion 1/2. To emulate population structure, the top 3 PCs of the empirical genetic relatedness matrix (GRM) were included as covariates. For all simulations, the target $\mu_{T,i}$ and surrogate $\mu_{S,i}$ regressions each included an intercept, age, sex, and 3 genetic PCs. Fixed effect regression coefficients were selected such that the proportion of total outcome-variation explained (PVE) by age and sex was 20%, and the PVE by genetic PCs was 5%. Initially, all simulations were performed in the presence ($\alpha_G \neq 0$) and absence ($\alpha_G = 0$) of an effect for genotype on the *surrogate* outcome. However, since $\alpha_G$ had no effect on the operating characteristics of SPRAY, including both size and power, only results for $\alpha_G = 0$ are reported here. For size simulations, genotype had no effect on the target outcome ($\beta_G = 0$). For power simulations, the effect of genotype was varied to achieve heritabilities between 0.1 and 1.0% ($\beta_G = 1/\sqrt{749}$ to $\beta_G = 1/\sqrt{74}$). Given genotype $g_i$ and covariates $\boldsymbol{x}_i$, the target and surrogate means were calculated as:

$$\mu_{T,i} = g_i\beta_G + \boldsymbol{x}_i'\boldsymbol{\beta}_X,$$
$$\mu_{S,i} = g_i\alpha_G + \boldsymbol{x}_i'\boldsymbol{\alpha}_X$$

The target and surrogate outcomes were generated by adding a residual vector to the subject-specific mean vector:

$$\begin{pmatrix} T_i \\ S_i \end{pmatrix} = \begin{pmatrix} \mu_{T,i} \\ \mu_{S,i} \end{pmatrix} + \begin{pmatrix} \epsilon_{T,i} \\ \epsilon_{S,i} \end{pmatrix} \tag{S3}$$

The target residual $\epsilon_{T,i}$ and surrogate proto-residual $\epsilon^*_{S,i}$ were drawn independently from a distribution with mean zero and variance one. The surrogate residual was then set to $\epsilon_{S,i} = \rho\epsilon_{T,i} + \sqrt{1-\rho^2}\epsilon^*_{S,i}$ such that $\mathbb{V}(\epsilon_{S,i}) = 1$ and $\mathbb{C}(\epsilon_{S,i}, \epsilon_{T,i}) = \rho$. For simulations under correct model specification, $\epsilon_{T,i}$ and $\epsilon^*_{S,i}$ were drawn from a standard normal distribution. For simulations with distributional misspecification, the residuals were from an exponential, log-normal, or $t_3$ distribution. In the case of the log-normal distribution, the residuals were chosen to have mean zero and unit variance on log-scale.

The number of complete cases was fixed at $n_0 = 10^3$. The numbers of subjects ($n_1$ and $n_2$) with target and surrogate missingness were varied to achieve specified proportions of target $\pi_T = n_1/(n_0 + n_1 + n_2)$ and surrogate $\pi_S = n_2/(n_0 + n_1 + n_2)$ missingness. For instance, when $\pi_T = 0.25$ and $\pi_S = 0.25$, the strata sizes were $n_0 = 10^3$ complete cases, $n_1 = 500$ subjects with target missingness, and $n_2 = 500$ subjects with surrogate missingness. Seven (target, surrogate) missingness patterns $(\pi_T, \pi_S)$ were considered: no missingness $(0.00, 0.00)$; unilateral target missingness $\{(0.25, 0.00), (0.50, 0.00), (0.75, 0.00)\}$; and bilateral outcome missingness $\{(0.25, 0.25), (0.50, 0.25), (0.25, 0.50)\}$. For each missingness pattern, the target-surrogate correlation $\rho$ spanned $\{0.00, 0.25, 0.50, 0.75\}$. When $\rho = 0$, the target and surrogate outcomes were in fact independent.

### 4 Estimation

#### 4.1 Target Parameters

**Supporting Table S1**: **Target parameter estimation and standard error calibration across $\boldsymbol{R = 5 \times 10^7}$ simulations in the presence of bilateral missingness**. The number of complete cases was $n_0 = 10^3$. The true regression coefficient ($\beta_G \approx 0.08$) was chosen such that the heritability of the target outcome was 0.5%. The true variance of the target outcome was $\Sigma_{TT} = 1.00$. The target missingness $\pi_T$, surrogate missingness $\pi_S$, and target-surrogate correlation $\rho$ were varied. The point estimate (EST) is the average across simulation replicates. The standard error is presented as the root mean square model-based standard error ($\text{SE}_M$), followed by the empirical standard error ($\text{SE}_E$) in parentheses, which is the standard deviation of the simulation point estimates.

| **Settings** | | | $\boldsymbol{\beta_G}$ | | $\boldsymbol{\Sigma_{TT}}$ | | $\boldsymbol{\rho}$ | |
|---|---|---|---|---|---|---|---|---|
| $\rho$ | $\pi_T$ | $\pi_S$ | EST | $\text{SE}_M$ ($\text{SE}_E$) | EST | $\text{SE}_M$ ($\text{SE}_E$) | EST | $\text{SE}_M$ ($\text{SE}_E$) |
| 0.00 | 0.25 | 0.25 | 0.08 | 0.04 (0.04) | 1.00 | 0.04 (0.04) | 0.00 | 0.03 (0.03) |
| 0.25 | 0.25 | 0.25 | 0.08 | 0.04 (0.04) | 1.00 | 0.04 (0.04) | 0.25 | 0.03 (0.03) |
| 0.50 | 0.25 | 0.25 | 0.08 | 0.04 (0.04) | 1.00 | 0.04 (0.04) | 0.50 | 0.03 (0.03) |
| 0.75 | 0.25 | 0.25 | 0.08 | 0.04 (0.04) | 1.00 | 0.04 (0.04) | 0.75 | 0.03 (0.03) |
| 0.00 | 0.25 | 0.50 | 0.08 | 0.03 (0.03) | 1.00 | 0.03 (0.03) | 0.00 | 0.03 (0.03) |
| 0.25 | 0.25 | 0.50 | 0.08 | 0.03 (0.03) | 1.00 | 0.03 (0.03) | 0.25 | 0.03 (0.03) |
| 0.50 | 0.25 | 0.50 | 0.08 | 0.03 (0.03) | 1.00 | 0.03 (0.03) | 0.50 | 0.03 (0.03) |
| 0.75 | 0.25 | 0.50 | 0.08 | 0.03 (0.03) | 1.00 | 0.02 (0.02) | 0.75 | 0.02 (0.02) |
| 0.00 | 0.50 | 0.25 | 0.08 | 0.04 (0.04) | 1.00 | 0.03 (0.03) | 0.00 | 0.03 (0.03) |
| 0.25 | 0.50 | 0.25 | 0.08 | 0.04 (0.04) | 1.00 | 0.03 (0.03) | 0.25 | 0.03 (0.03) |
| 0.50 | 0.50 | 0.25 | 0.08 | 0.03 (0.03) | 1.00 | 0.03 (0.03) | 0.50 | 0.03 (0.03) |
| 0.75 | 0.50 | 0.25 | 0.08 | 0.03 (0.03) | 1.00 | 0.03 (0.03) | 0.75 | 0.02 (0.02) |

#### 4.2 Surrogate Parameters

**Supporting Table S2**: **Surrogate parameter estimation and standard error calibration across $\boldsymbol{R = 5 \times 10^7}$ simulations**. The number of complete cases was $n_0 = 10^3$. The true regression coefficient ($\alpha_G \approx 0.08$) was chosen such that the heritability of the surrogate outcome was 0.5%. The surrogate missingness $\pi_S = 0$ was fixed while the target missingness $\pi_T$ and target-surrogate correlation $\rho$ were varied. The point estimate (EST) is the average across simulation replicates. The standard error is presented as the root mean square model-based standard error ($\text{SE}_M$), followed by the empirical standard error ($\text{SE}_E$) in parentheses, which is the standard deviation of the simulation point estimates.

| **Settings** | | | $\boldsymbol{\alpha_G}$ | | $\boldsymbol{\Sigma_{SS}}$ | |
|---|---|---|---|---|---|---|
| $\rho$ | $\pi_T$ | $\pi_S$ | EST | $\text{SE}_M$ ($\text{SE}_E$) | EST | $\text{SE}_M$ ($\text{SE}_E$) |
| 0.00 | 0.00 | 0.00 | 0.08 | 0.05 (0.05) | 0.99 | 0.04 (0.04) |
| 0.25 | 0.00 | 0.00 | 0.08 | 0.05 (0.05) | 0.99 | 0.04 (0.04) |
| 0.50 | 0.00 | 0.00 | 0.08 | 0.05 (0.05) | 1.00 | 0.04 (0.04) |
| 0.75 | 0.00 | 0.00 | 0.08 | 0.05 (0.05) | 0.99 | 0.04 (0.05) |
| 0.00 | 0.25 | 0.00 | 0.08 | 0.04 (0.04) | 1.00 | 0.04 (0.04) |
| 0.25 | 0.25 | 0.00 | 0.08 | 0.04 (0.04) | 1.00 | 0.04 (0.04) |
| 0.50 | 0.25 | 0.00 | 0.08 | 0.04 (0.04) | 0.99 | 0.04 (0.04) |
| 0.75 | 0.25 | 0.00 | 0.08 | 0.04 (0.04) | 1.00 | 0.04 (0.04) |
| 0.00 | 0.50 | 0.00 | 0.08 | 0.04 (0.04) | 1.00 | 0.03 (0.03) |
| 0.25 | 0.50 | 0.00 | 0.08 | 0.04 (0.04) | 1.00 | 0.03 (0.03) |
| 0.50 | 0.50 | 0.00 | 0.08 | 0.04 (0.04) | 1.00 | 0.03 (0.03) |
| 0.75 | 0.50 | 0.00 | 0.08 | 0.04 (0.04) | 1.00 | 0.03 (0.03) |
| 0.00 | 0.75 | 0.00 | 0.08 | 0.03 (0.03) | 1.00 | 0.02 (0.02) |
| 0.25 | 0.75 | 0.00 | 0.08 | 0.03 (0.03) | 1.00 | 0.02 (0.02) |
| 0.50 | 0.75 | 0.00 | 0.08 | 0.03 (0.03) | 1.00 | 0.02 (0.02) |
| 0.75 | 0.75 | 0.00 | 0.08 | 0.03 (0.03) | 1.00 | 0.02 (0.02) |
| 0.00 | 0.25 | 0.25 | 0.08 | 0.04 (0.04) | 1.00 | 0.04 (0.04) |
| 0.25 | 0.25 | 0.25 | 0.08 | 0.04 (0.04) | 1.00 | 0.04 (0.04) |
| 0.50 | 0.25 | 0.25 | 0.08 | 0.04 (0.04) | 1.00 | 0.04 (0.04) |
| 0.75 | 0.25 | 0.25 | 0.08 | 0.04 (0.04) | 1.00 | 0.04 (0.04) |
| 0.00 | 0.25 | 0.50 | 0.08 | 0.04 (0.04) | 1.00 | 0.03 (0.03) |
| 0.25 | 0.25 | 0.50 | 0.08 | 0.04 (0.04) | 1.00 | 0.03 (0.03) |
| 0.50 | 0.25 | 0.50 | 0.08 | 0.03 (0.03) | 1.00 | 0.03 (0.03) |
| 0.75 | 0.25 | 0.50 | 0.08 | 0.03 (0.03) | 1.00 | 0.03 (0.03) |
| 0.00 | 0.50 | 0.25 | 0.08 | 0.03 (0.03) | 1.00 | 0.03 (0.03) |
| 0.25 | 0.50 | 0.25 | 0.08 | 0.03 (0.03) | 1.00 | 0.03 (0.03) |
| 0.50 | 0.50 | 0.25 | 0.08 | 0.03 (0.03) | 1.00 | 0.03 (0.03) |
| 0.75 | 0.50 | 0.25 | 0.08 | 0.03 (0.03) | 1.00 | 0.03 (0.02) |

#### 4.3 Model Misspecification

The model misspecification simulations focus on assessing the accuracy and calibration of estimates for the genetic effect $\beta_G$, which is the parameter of primary interest. Phenotypes were generated via equation (S3). The residual distributions were bivariate normal (reference), exponential, log-normal, and Student with 3 degrees of freedom ($t_3$). The target and surrogate outcomes were each INT transformed prior to estimation.

**Supporting Table S3**: **Unbiased estimation of the genetic effect $\boldsymbol{\beta_G}$ across $\boldsymbol{R = 5 \times 10^7}$ simulations under model misspecification**. The number of complete cases was $n_0 = 10^3$. The true regression coefficient ($\beta_G \approx 0.08$) was chosen such that the heritability of the surrogate outcome was 0.5%. The surrogate missingness $\pi_S = 0$ was fixed while the target missingness $\pi_T$ and target-surrogate correlation $\rho$ were varied. Bias is the average discrepancy between the estimated and true $\beta_G$. The standard error is presented as the root mean square model-based standard error ($\text{SE}_M$), followed by the empirical standard error ($\text{SE}_E$) in parentheses, which is the standard deviation of the simulation point estimates.

| **Settings** | | **Normal** | | **Exponential** | | **Log-Normal** | | **Student $t_3$** | |
|---|---|---|---|---|---|---|---|---|---|
| $\rho$ | $\pi_T$ | Bias | $\text{SE}_M$ ($\text{SE}_E$) | Bias | $\text{SE}_M$ ($\text{SE}_E$) | Bias | $\text{SE}_M$ ($\text{SE}_E$) | Bias | $\text{SE}_M$ ($\text{SE}_E$) |
| 0.00 | 0.00 | 0.000 | 0.032 (0.032) | -0.000 | 0.032 (0.032) | 0.000 | 0.032 (0.032) | 0.000 | 0.032 (0.032) |
| 0.25 | 0.00 | 0.000 | 0.032 (0.032) | 0.000 | 0.032 (0.032) | 0.000 | 0.032 (0.032) | -0.000 | 0.032 (0.032) |
| 0.50 | 0.00 | 0.000 | 0.032 (0.032) | 0.000 | 0.032 (0.032) | 0.000 | 0.032 (0.032) | 0.000 | 0.032 (0.032) |
| 0.75 | 0.00 | 0.000 | 0.032 (0.032) | -0.000 | 0.032 (0.032) | 0.000 | 0.032 (0.032) | -0.000 | 0.032 (0.032) |
| 0.00 | 0.25 | -0.000 | 0.032 (0.032) | -0.000 | 0.032 (0.032) | -0.000 | 0.032 (0.032) | -0.000 | 0.032 (0.032) |
| 0.25 | 0.25 | 0.000 | 0.032 (0.032) | 0.000 | 0.032 (0.032) | -0.000 | 0.032 (0.032) | 0.000 | 0.032 (0.032) |
| 0.50 | 0.25 | -0.000 | 0.031 (0.031) | -0.000 | 0.032 (0.032) | 0.000 | 0.032 (0.032) | 0.000 | 0.032 (0.032) |
| 0.75 | 0.25 | -0.000 | 0.030 (0.030) | -0.000 | 0.032 (0.032) | -0.000 | 0.032 (0.032) | 0.000 | 0.032 (0.032) |
| 0.00 | 0.50 | 0.000 | 0.032 (0.032) | -0.000 | 0.032 (0.032) | 0.000 | 0.032 (0.032) | 0.000 | 0.032 (0.032) |
| 0.25 | 0.50 | -0.000 | 0.032 (0.032) | 0.000 | 0.032 (0.032) | 0.000 | 0.032 (0.032) | 0.000 | 0.032 (0.032) |
| 0.50 | 0.50 | -0.000 | 0.030 (0.030) | -0.000 | 0.032 (0.032) | -0.000 | 0.032 (0.032) | 0.000 | 0.032 (0.032) |
| 0.75 | 0.50 | -0.000 | 0.027 (0.027) | -0.000 | 0.032 (0.032) | -0.000 | 0.032 (0.032) | 0.000 | 0.032 (0.032) |
| 0.00 | 0.75 | -0.000 | 0.032 (0.032) | 0.000 | 0.032 (0.032) | -0.000 | 0.032 (0.032) | 0.000 | 0.032 (0.032) |
| 0.25 | 0.75 | -0.000 | 0.031 (0.031) | -0.000 | 0.032 (0.032) | 0.000 | 0.032 (0.032) | 0.000 | 0.032 (0.032) |
| 0.50 | 0.75 | -0.000 | 0.029 (0.029) | 0.000 | 0.032 (0.032) | -0.000 | 0.032 (0.032) | 0.000 | 0.032 (0.032) |
| 0.75 | 0.75 | 0.000 | 0.024 (0.024) | -0.000 | 0.032 (0.032) | 0.000 | 0.032 (0.032) | -0.000 | 0.032 (0.032) |

### 5 Type I Error

#### 5.1 Unilateral Missingness

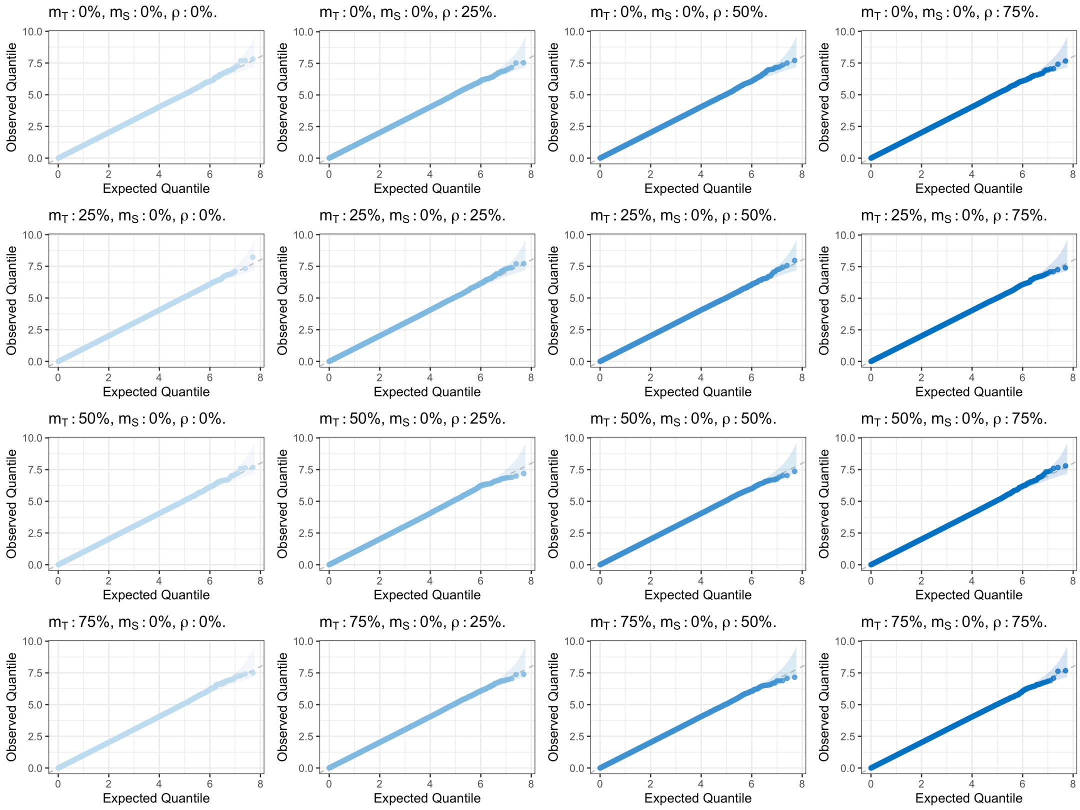

**Supporting Figure S1**: **Quantile-quantile plots comparing the observed distribution of p-values with the standard uniform distribution across $\boldsymbol{R = 5 \times 10^7}$ simulation replicates in the presence of unilateral missingness**. The number of complete cases is $n_0 = 10^3$. $\rho$ is the target-surrogate correlation; $m_T$ is missingness in the target outcome; $m_S$ is missingness in the surrogate outcome.

#### 5.2 Bilateral Missingness

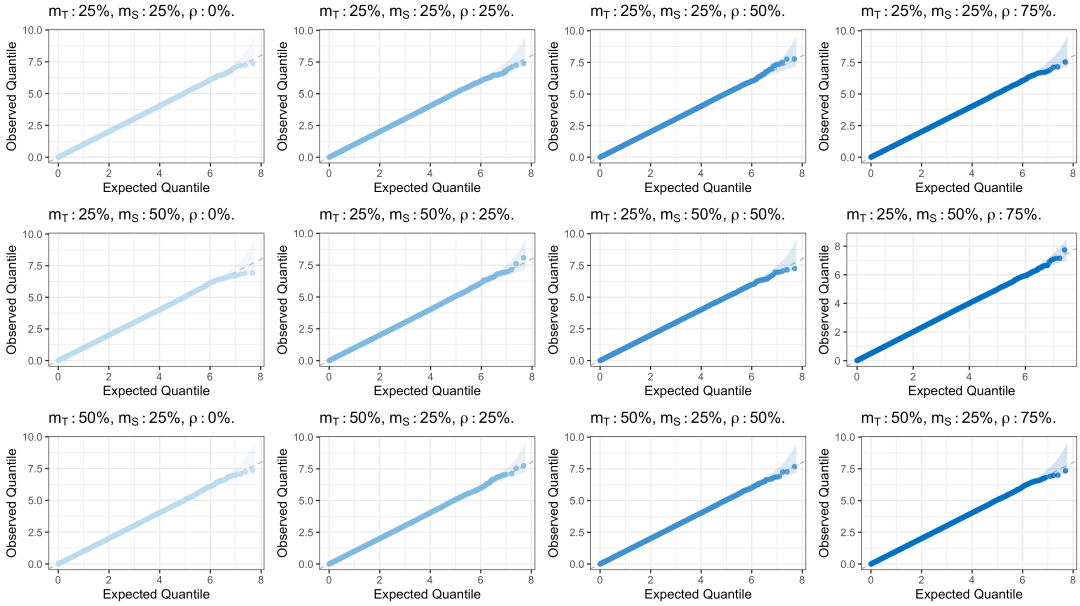

**Supporting Figure S2**: **Quantile-quantile plots comparing the observed distribution of p-values with the standard uniform distribution across $\boldsymbol{R = 5 \times 10^7}$ simulation replicates in the presence of bilateral missingness**. The number of complete cases is $n_0 = 10^3$. $\rho$ is the target-surrogate correlation; $m_T$ is missingness in the target outcome; $m_S$ is missingness in the surrogate outcome.

**Supporting Table S4**: **Empirical type I error and power of the Spray Wald test across $\boldsymbol{R} = 5 \times 10^7$ simulation replicates in the presence of bilateral missingness**. The number of complete cases was $n_0 = 10^3$. The surrogate missingness was fixed at $\pi_S = 0$. For type I error, $\beta_G = 0$ while for power $\beta_G$ was selected to explain 0.5% of variation in the target outcome. The target missingness $\pi_T$, surrogate missingness $\pi_S$, and target-surrogate correlation $\rho$ were varied. Prob refers to the rejection probability at a target type I error of 5% and NCP is the non-centrality parameter of the Wald test.

| **Settings** | | | **Type I Error** | | **Power** | |
|---|---|---|---|---|---|---|
| $\rho$ | $\pi_T$ | $\pi_S$ | Prob (%) | NCP | Prob (%) | NCP |
| 0.00 | 0.25 | 0.25 | 5.08 | 1.01 | 88.00 | 10.86 |
| 0.25 | 0.25 | 0.25 | 5.08 | 1.01 | 88.40 | 10.99 |
| 0.50 | 0.25 | 0.25 | 5.07 | 1.01 | 89.79 | 11.46 |
| 0.75 | 0.25 | 0.25 | 5.07 | 1.01 | 91.83 | 12.33 |
| 0.00 | 0.25 | 0.50 | 5.04 | 1.00 | 99.32 | 20.77 |
| 0.25 | 0.25 | 0.50 | 5.03 | 1.00 | 99.35 | 20.89 |
| 0.50 | 0.25 | 0.50 | 5.04 | 1.00 | 99.49 | 21.72 |
| 0.75 | 0.25 | 0.50 | 5.04 | 1.00 | 99.69 | 23.29 |
| 0.00 | 0.50 | 0.25 | 5.06 | 1.01 | 95.10 | 14.15 |
| 0.25 | 0.50 | 0.25 | 5.07 | 1.01 | 95.57 | 14.44 |
| 0.50 | 0.50 | 0.25 | 5.06 | 1.00 | 96.65 | 15.48 |
| 0.75 | 0.50 | 0.25 | 5.05 | 1.00 | 98.46 | 18.09 |

#### 5.3 Model Misspecification

##### 5.3.1 Exponential

**Supporting Table S5**: **Empirical type I error and power of the Spray Wald test across $\boldsymbol{R} = \mathbf{10^7}$ simulation replicates under an exponential data generating process**. The number of complete cases was $n_0 = 10^3$. The surrogate missingness $\pi_S = 0$ was fixed while the target missingness $\pi_T$ and target-surrogate correlation $\rho$ were varied. For type I error, $\beta_G = 0$ while for power $\beta_G$ was selected to explain 0.5% of variation in the target outcome. Prob refers to the rejection probability at a target type I error of 5% and NCP is the non-centrality parameter of the Wald test. The target and surrogate outcomes were each rank-normalized prior to estimation.

| **Settings** | | **Type I Error** | | **Power** | |
|---|---|---|---|---|---|
| $\rho$ | $\pi_T$ | Prob (%) | NCP | Prob (%) | NCP |
| 0.00 | 0.00 | 5.01 | 1.00 | 71.32 | 7.42 |
| 0.25 | 0.00 | 5.01 | 1.00 | 71.17 | 7.39 |
| 0.50 | 0.00 | 5.01 | 1.00 | 71.00 | 7.40 |
| 0.75 | 0.00 | 5.02 | 1.00 | 71.15 | 7.41 |
| 0.00 | 0.25 | 5.02 | 1.00 | 70.92 | 7.39 |
| 0.25 | 0.25 | 5.03 | 1.00 | 71.86 | 7.51 |
| 0.50 | 0.25 | 5.02 | 1.00 | 74.03 | 7.85 |
| 0.75 | 0.25 | 5.02 | 1.00 | 77.65 | 8.46 |
| 0.00 | 0.50 | 5.03 | 1.00 | 71.49 | 7.48 |
| 0.25 | 0.50 | 5.03 | 1.00 | 73.01 | 7.69 |
| 0.50 | 0.50 | 5.02 | 1.00 | 77.20 | 8.37 |
| 0.75 | 0.50 | 5.03 | 1.00 | 84.88 | 9.98 |
| 0.00 | 0.75 | 5.04 | 1.00 | 71.63 | 7.47 |
| 0.25 | 0.75 | 5.04 | 1.00 | 73.43 | 7.77 |
| 0.50 | 0.75 | 5.04 | 1.00 | 80.03 | 8.92 |
| 0.75 | 0.75 | 5.04 | 1.00 | 91.48 | 12.17 |

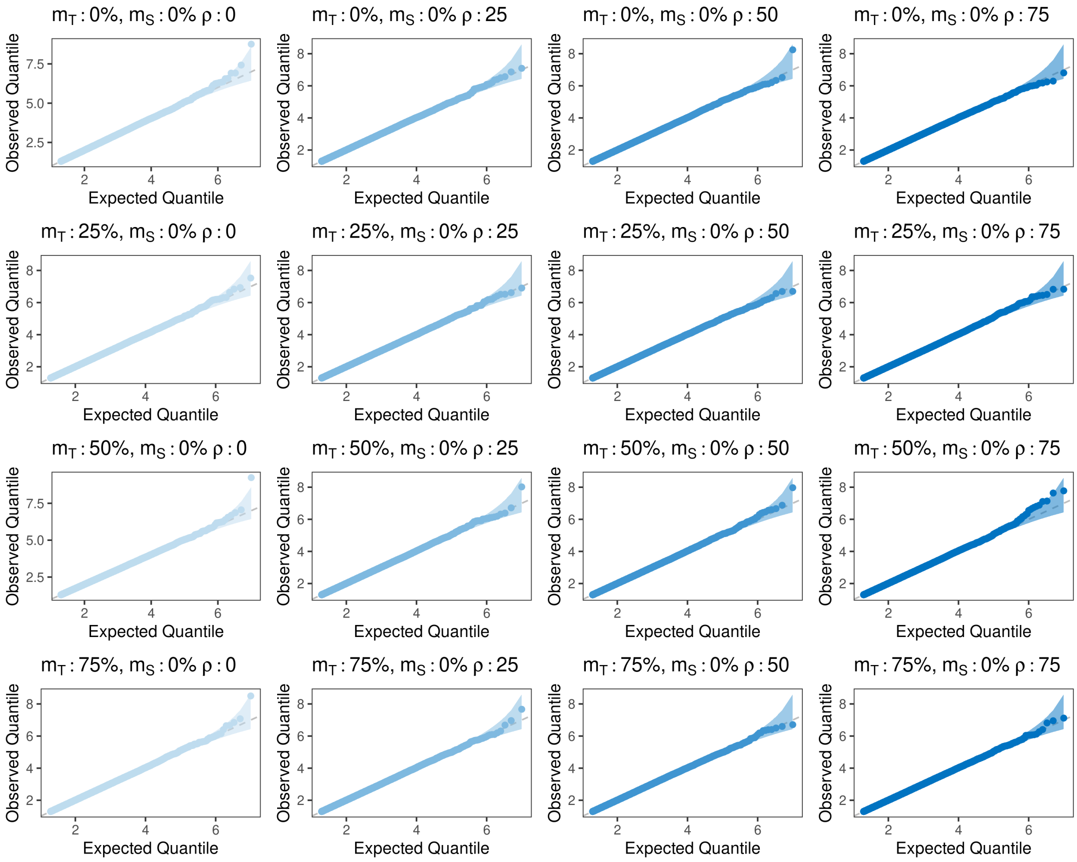

**Supporting Figure S3**: **Quantile-quantile plots comparing the observed distribution of p-values with the standard uniform distribution across $\boldsymbol{R} = \mathbf{10^7}$ simulation replicates under an exponential data generating process**. The number of complete cases is $n_0 = 10^3$. $\rho$ is the target-surrogate correlation; $m_T$ is missingness in the target outcome; $m_S$ is missingness in the surrogate outcome.

##### 5.3.2 Log-Normal

**Supporting Table S6**: **Empirical type I error and power of the Spray Wald test across $\boldsymbol{R} = \mathbf{10^7}$ simulation replicates under a log-normal generating process**. The number of complete cases was $n_0 = 10^3$. The surrogate missingness $\pi_S = 0$ was fixed while the target missingness $\pi_T$ and target-surrogate correlation $\rho$ were varied. For type I error, $\beta_G = 0$ while for power $\beta_G$ was selected to explain 0.5% of variation in the target outcome. Prob refers to the rejection probability at a target type I error of 5% and NCP is the non-centrality parameter of the Wald test. The target and surrogate outcomes were each rank-normalized prior to estimation.

| **Settings** | | **Type I Error** | | **Power** | |
|---|---|---|---|---|---|
| $\rho$ | $\pi_T$ | Prob (%) | NCP | Prob (%) | NCP |
| 0.00 | 0.00 | 5.01 | 1.00 | 22.23 | 2.45 |
| 0.25 | 0.00 | 5.02 | 1.00 | 22.52 | 2.45 |
| 0.50 | 0.00 | 5.01 | 1.00 | 22.49 | 2.47 |
| 0.75 | 0.00 | 5.02 | 1.00 | 22.09 | 2.44 |
| 0.00 | 0.25 | 5.02 | 1.00 | 22.39 | 2.45 |
| 0.25 | 0.25 | 5.03 | 1.00 | 22.75 | 2.48 |
| 0.50 | 0.25 | 5.02 | 1.00 | 22.92 | 2.50 |
| 0.75 | 0.25 | 5.03 | 1.00 | 24.52 | 2.63 |
| 0.00 | 0.50 | 5.02 | 1.00 | 22.46 | 2.46 |
| 0.25 | 0.50 | 5.02 | 1.00 | 22.67 | 2.48 |
| 0.50 | 0.50 | 5.02 | 1.00 | 23.88 | 2.58 |
| 0.75 | 0.50 | 5.02 | 1.00 | 27.12 | 2.84 |
| 0.00 | 0.75 | 5.04 | 1.00 | 22.69 | 2.48 |
| 0.25 | 0.75 | 5.04 | 1.00 | 23.01 | 2.50 |
| 0.50 | 0.75 | 5.02 | 1.00 | 24.95 | 2.67 |
| 0.75 | 0.75 | 5.04 | 1.00 | 30.73 | 3.16 |

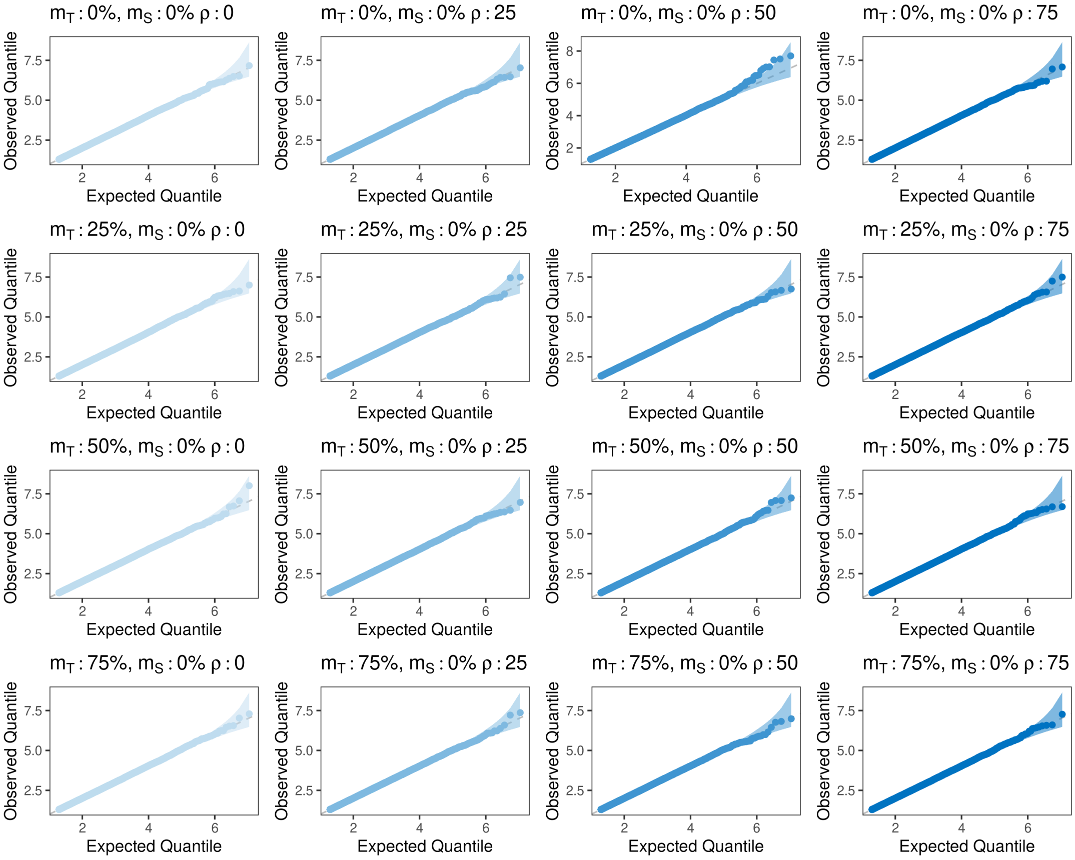

**Supporting Figure S4**: **Quantile-quantile plots comparing the observed distribution of p-values with the standard uniform distribution across $\boldsymbol{R} = \mathbf{10^7}$ simulation replicates under a log-normal data generating process**. The number of complete cases is $n_0 = 10^3$. $\rho$ is the target-surrogate correlation; $m_T$ is missingness in the target outcome; $m_S$ is missingness in the surrogate outcome.

##### 5.3.3 Student

**Supporting Table S7**: **Empirical type I error and power of the Spray Wald test across $\boldsymbol{R = 10^7}$ simulation replicates under a Student $\boldsymbol{t_3}$ generating process**. The number of complete cases was $n_0 = 10^3$. The surrogate missingness $\pi_S = 0$ was fixed while the target missingness $\pi_T$ and target-surrogate correlation $\rho$ were varied. For type I error, $\beta_G = 0$ while for power $\beta_G$ was selected to explain 0.5% of variation in the target outcome. Prob refers to the rejection probability at a target type I error of 5% and NCP is the non-centrality parameter of the Wald test. The target and surrogate outcomes were each rank-normalized prior to estimation.

| **Settings** | | **Type I Error** | | **Power** | |
|---|---|---|---|---|---|
| $\rho$ | $\pi_T$ | Prob (%) | NCP | Prob (%) | NCP |
| 0.00 | 0.00 | 5.02 | 1.00 | 32.61 | 3.28 |
| 0.25 | 0.00 | 5.01 | 1.00 | 32.50 | 3.27 |
| 0.50 | 0.00 | 5.01 | 1.00 | 32.27 | 3.27 |
| 0.75 | 0.00 | 5.01 | 1.00 | 32.32 | 3.26 |
| 0.00 | 0.25 | 5.04 | 1.00 | 32.55 | 3.28 |
| 0.25 | 0.25 | 5.02 | 1.00 | 32.66 | 3.29 |
| 0.50 | 0.25 | 5.03 | 1.00 | 34.28 | 3.42 |
| 0.75 | 0.25 | 5.02 | 1.00 | 36.47 | 3.61 |
| 0.00 | 0.50 | 5.03 | 1.00 | 32.83 | 3.31 |
| 0.25 | 0.50 | 5.04 | 1.00 | 32.96 | 3.32 |
| 0.50 | 0.50 | 5.03 | 1.00 | 36.49 | 3.60 |
| 0.75 | 0.50 | 5.03 | 1.00 | 42.23 | 4.12 |
| 0.00 | 0.75 | 5.02 | 1.00 | 32.71 | 3.30 |
| 0.25 | 0.75 | 5.04 | 1.00 | 34.22 | 3.41 |
| 0.50 | 0.75 | 5.04 | 1.00 | 38.80 | 3.82 |
| 0.75 | 0.75 | 5.03 | 1.00 | 50.72 | 4.92 |

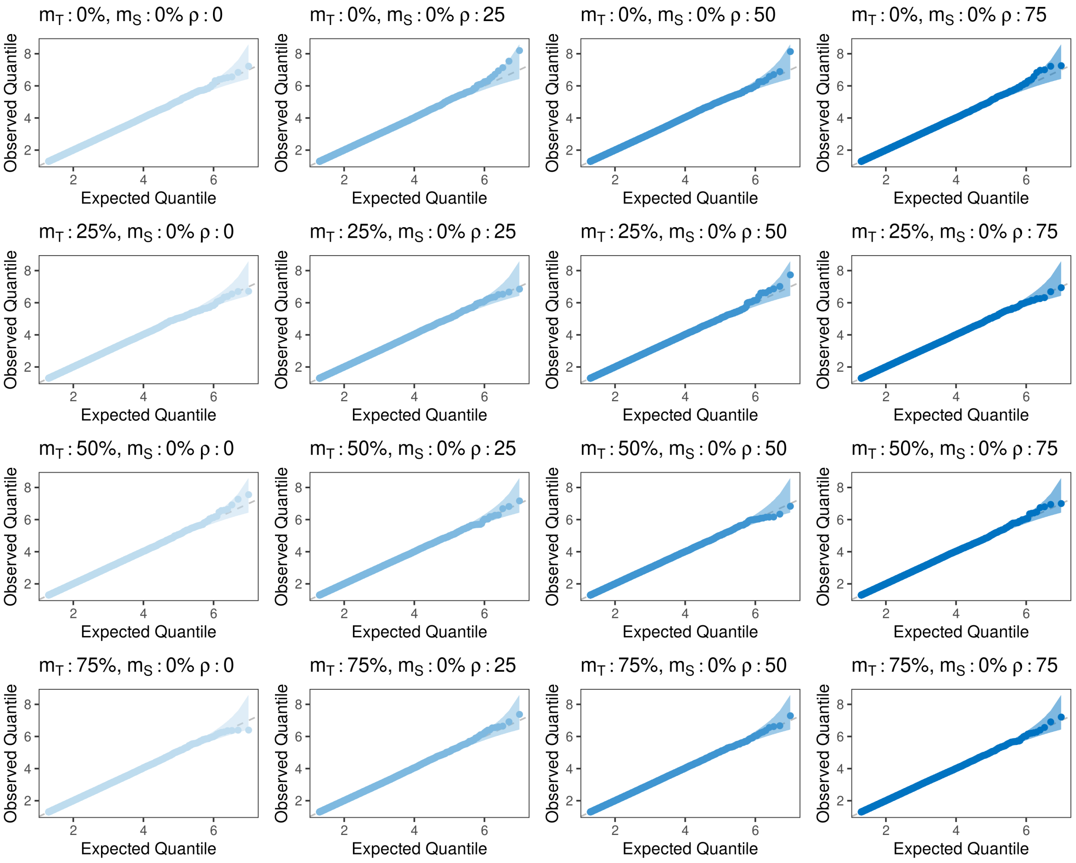

**Supporting Figure S5**: **Quantile-quantile plots comparing the observed distribution of p-values with the standard uniform distribution across $\boldsymbol{R = 10^7}$ simulation replicates under a Student $\boldsymbol{t_3}$ generating process.** The number of complete cases is $n_0 = 10^3$. $\rho$ is the target-surrogate correlation; $m_T$ is missingness in the target outcome; $m_S$ is missingness in the surrogate outcome.

#### 5.4 Large Sample

**Supporting Table S8: Empirical type I error and power of the Spray Wald test across $\boldsymbol{R} = 5 \times 10^6$ simulation replicates at increasing sample size**. The target missingness $\pi_T$, surrogate missingness $\pi_S$, and target-surrogate correlation $\rho$ were all varied. The true $\beta_G = 0$. Prob refers to the rejection probability at a target type I error of 5% and NCP is the non-centrality parameter of the Wald test.

| **Settings** | | | $n = 5000$ | | $n = 10000$ | | $n = 20000$ | |
|---|---|---|---|---|---|---|---|---|
| $\rho$ | $\pi_T$ | $\pi_S$ | Prob (%) | NCP | Prob (%) | NCP | Prob (%) | NCP |
| 0.00 | 0.00 | 0.00 | 4.999 | 1.000 | 4.993 | 1.000 | 5.004 | 1.000 |
| 0.25 | 0.00 | 0.00 | 5.020 | 1.001 | 4.997 | 1.000 | 5.000 | 1.000 |
| 0.50 | 0.00 | 0.00 | 5.004 | 1.000 | 4.994 | 1.000 | 5.001 | 1.000 |
| 0.75 | 0.00 | 0.00 | 4.989 | 0.999 | 5.016 | 1.000 | 4.994 | 0.999 |
| 0.00 | 0.25 | 0.00 | 5.016 | 1.001 | 5.001 | 1.000 | 4.994 | 1.000 |
| 0.25 | 0.25 | 0.00 | 5.002 | 1.000 | 5.003 | 1.001 | 5.007 | 1.000 |
| 0.50 | 0.25 | 0.00 | 5.006 | 1.001 | 4.993 | 0.999 | 5.005 | 1.000 |
| 0.75 | 0.25 | 0.00 | 5.021 | 1.001 | 5.002 | 0.999 | 5.006 | 1.001 |
| 0.00 | 0.50 | 0.00 | 5.007 | 1.000 | 5.015 | 1.001 | 4.985 | 0.999 |
| 0.25 | 0.50 | 0.00 | 5.017 | 1.001 | 5.012 | 1.001 | 5.008 | 1.001 |
| 0.50 | 0.50 | 0.00 | 5.015 | 1.001 | 4.997 | 1.000 | 5.003 | 1.000 |
| 0.75 | 0.50 | 0.00 | 5.012 | 1.001 | 4.986 | 0.999 | 5.001 | 1.000 |
| 0.00 | 0.75 | 0.00 | 5.009 | 1.000 | 5.009 | 1.001 | 5.006 | 1.000 |
| 0.25 | 0.75 | 0.00 | 4.990 | 1.000 | 5.004 | 1.001 | 5.003 | 1.000 |
| 0.50 | 0.75 | 0.00 | 5.019 | 1.001 | 5.012 | 1.001 | 4.981 | 0.999 |
| 0.75 | 0.75 | 0.00 | 5.017 | 1.001 | 5.003 | 1.001 | 4.983 | 0.999 |
| 0.00 | 0.25 | 0.25 | 5.029 | 1.001 | 5.011 | 1.000 | 5.005 | 1.000 |
| 0.25 | 0.25 | 0.25 | 5.004 | 1.001 | 5.001 | 1.000 | 5.000 | 0.999 |
| 0.50 | 0.25 | 0.25 | 5.009 | 1.001 | 5.005 | 1.001 | 5.007 | 1.001 |
| 0.75 | 0.25 | 0.25 | 5.018 | 1.001 | 5.008 | 1.001 | 5.000 | 1.000 |
| 0.00 | 0.50 | 0.25 | 5.020 | 1.002 | 5.012 | 1.001 | 4.996 | 1.000 |
| 0.25 | 0.50 | 0.25 | 5.005 | 1.001 | 5.006 | 1.001 | 5.008 | 1.001 |
| 0.50 | 0.50 | 0.25 | 4.993 | 1.000 | 5.002 | 1.000 | 4.989 | 1.000 |
| 0.75 | 0.50 | 0.25 | 5.009 | 1.001 | 5.013 | 1.001 | 4.994 | 0.999 |
| 0.00 | 0.25 | 0.50 | 5.005 | 1.000 | 5.004 | 1.000 | 5.008 | 1.000 |
| 0.25 | 0.25 | 0.50 | 4.999 | 1.000 | 4.985 | 0.999 | 4.994 | 1.000 |
| 0.50 | 0.25 | 0.50 | 5.012 | 1.000 | 5.010 | 1.001 | 5.003 | 1.000 |
| 0.75 | 0.25 | 0.50 | 5.003 | 1.001 | 5.002 | 1.000 | 4.999 | 1.000 |

### 6 Power

#### 6.1 Bilateral Missingness

Note that these power curves saturate faster than those in the main text because the simulations include more subjects with observed target outcomes overall. For example, in the case of $\pi_T = 25\%$ and $\pi_S = 50\%$, the number of complete cases is $n_0 = 1000$, the number of subjects with target missingness is $n_1 = 1000$, and the number of subjects with surrogate missingness is $n_2 = 2000$. Therefore, the total number of subjects with observed target outcomes is $n_0 + n_2 = 3000$.

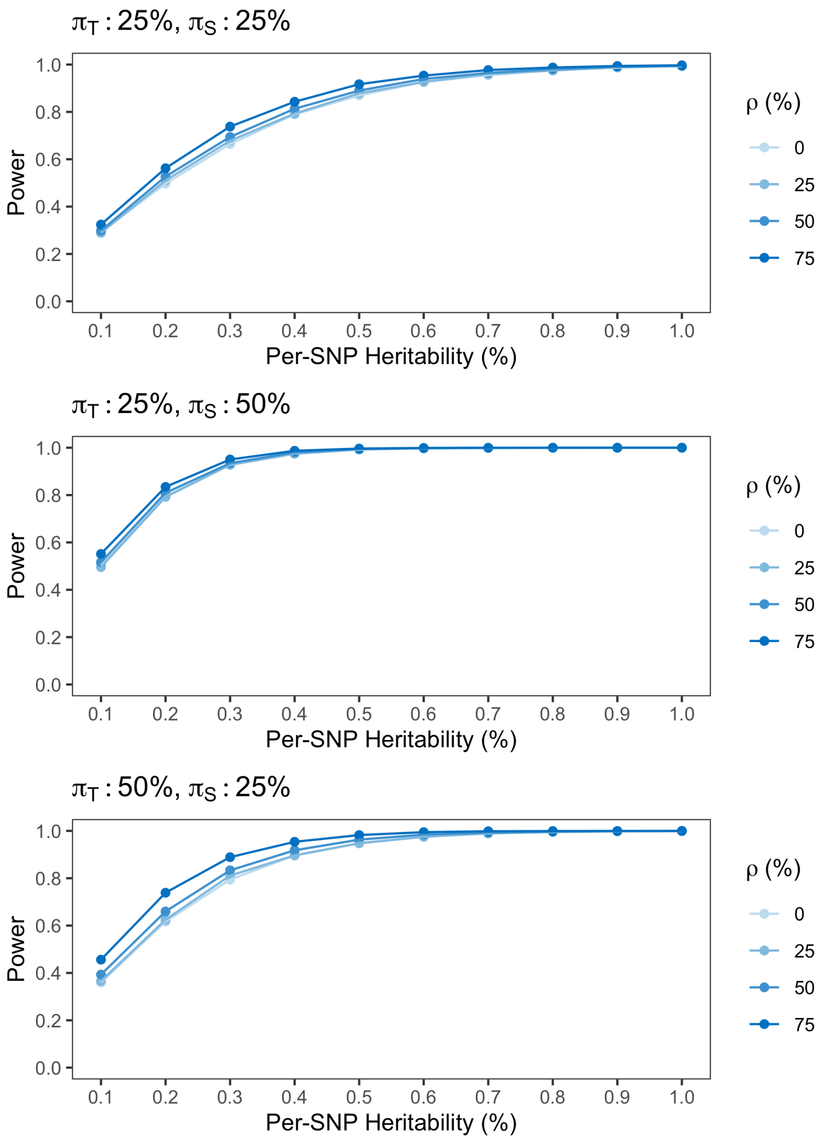

**Supporting Figure S6**: **Power curves for the Spray Wald test in the presence of bilateral missingness**. The number of complete cases was $n_0 = 10^3$, and the type I error was $\alpha = 0.05$. Each point on the curve corresponds to $R = 5 \times 10^5$ simulation replicates. $\rho$ is the target-surrogate correlation; $\pi_T$ is missingness in the target outcome; $\pi_S$ is missingness in the surrogate outcome.

#### 6.2 Model Misspecification

##### 6.2.1 Exponential

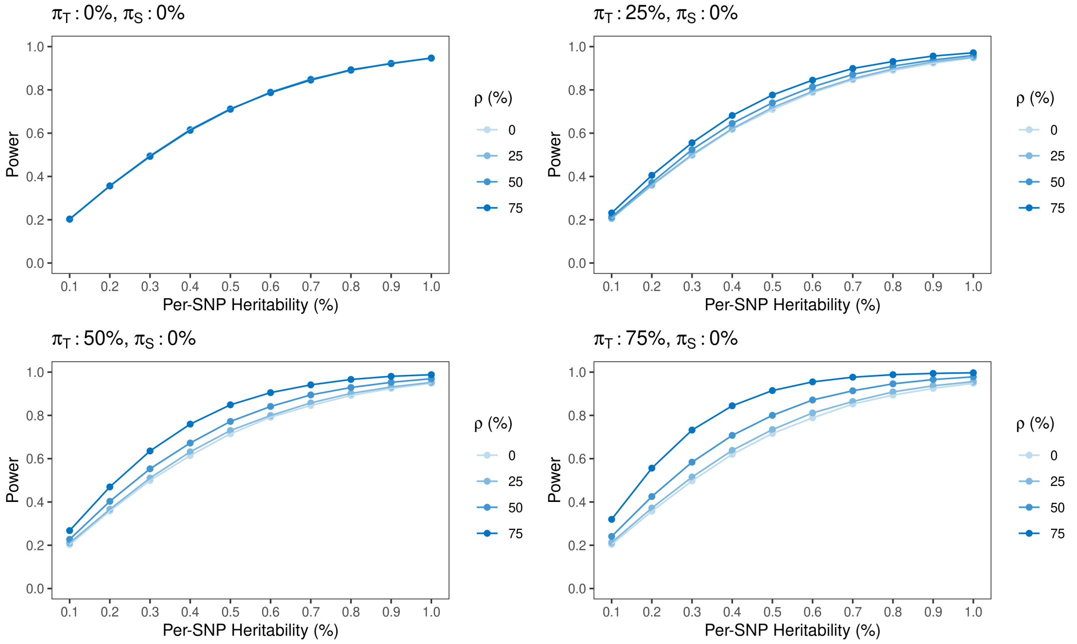

**Supporting Figure S7**: **Power curves for the Spray Wald test under an exponential generating process**. The number of complete cases was $n_0 = 10^3$, and the type I error was $\alpha = 0.05$. Each point on the curve corresponds to $R = 5 \times 10^5$ simulation replicates. $\rho$ is the target-surrogate correlation; $\pi_T$ is missingness in the target outcome; $\pi_S$ is missingness in the surrogate outcome.

##### 6.2.2 Log-Normal

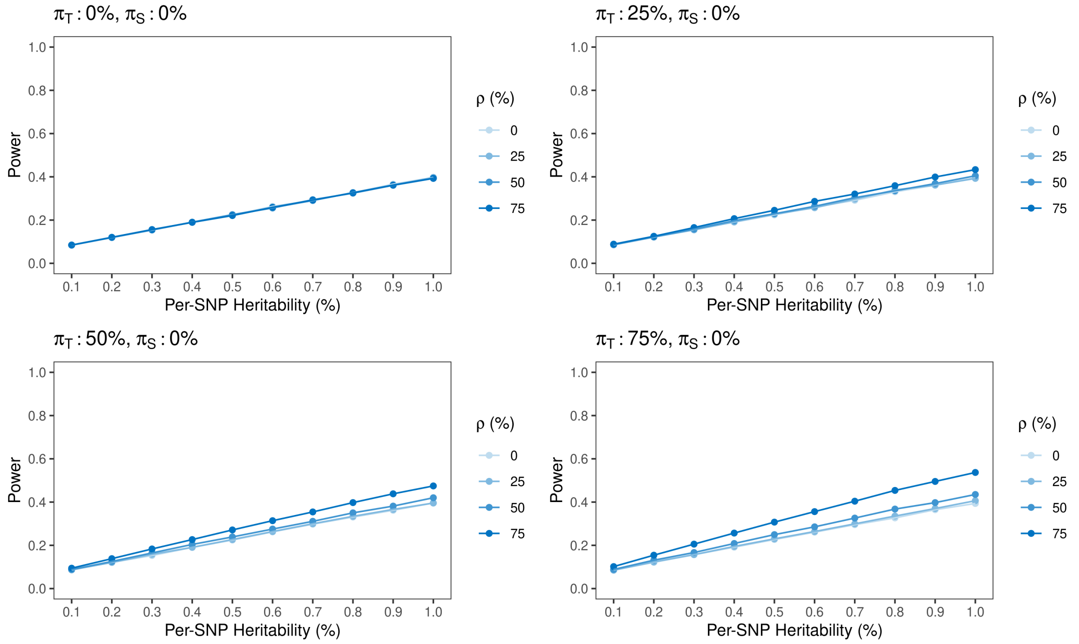

**Supporting Figure S8**: **Power curves for the Spray Wald test under a log-normal generating process**. The number of complete cases was $n_0 = 10^3$, and the type I error was $\alpha = 0.05$. Each point on the curve corresponds to $R = 5 \times 10^5$ simulation replicates. $\rho$ is the target-surrogate correlation; $\pi_T$ is missingness in the target outcome; $\pi_S$ is missingness in the surrogate outcome.

##### 6.2.3 Student

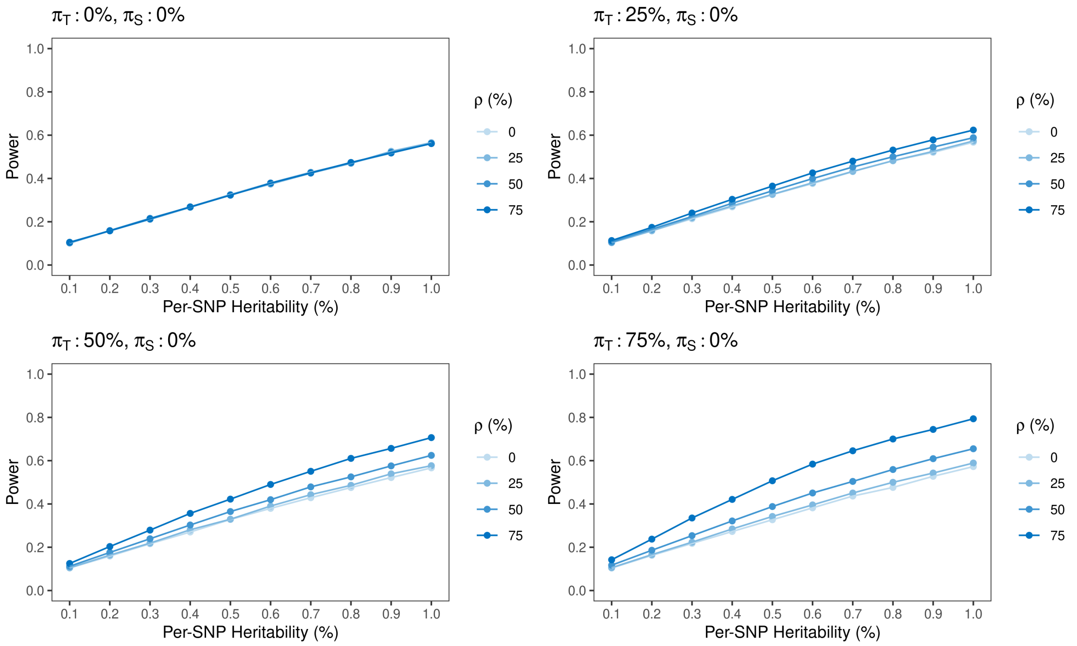

**Supporting Figure S9**: **Power curves for the Spray Wald test under a a Student $t_3$ generating process**. The number of complete cases was $n_0 = 10^3$, and the type I error was $\alpha = 0.05$. Each point on the curve corresponds to $R = 5 \times 10^5$ simulation replicates. $\rho$ is the target-surrogate correlation; $\pi_T$ is missingness in the target outcome; $\pi_S$ is missingness in the surrogate outcome.

### 7 Empirical Relative Efficiency

#### 7.1 Bilateral Missingness

**Supporting Table S9**: **Empirical relative efficiency of the Spray estimator to the marginal estimator of $\boldsymbol{\beta_G}$ test across $\boldsymbol{R = 5 \times 10^7}$ simulation replicates in the presence of bilateral missingness**. The number of complete cases was $n_0 = 10^3$. The true regression coefficients ($\beta_G \approx 0.08$) were chosen such that the heritability of each outcome was 0.5%. The true variances of the target and surrogate outcomes were $\Sigma_{TT} = \Sigma_{SS} = 1.00$. The target missingness $\pi_T$, surrogate missingness $\pi_S$, and target-surrogate correlation $\rho$ were varied. Variance refers to the empirical variance of the corresponding estimator across simulation replicates. The empirical RE is the ratio of the variance of $\hat{\beta}_G^{\text{Spray}}$ to that of $\hat{\beta}_G^{\text{Marginal}}$. The theoretical RE was obtained from (11).

| **Settings** | | | **Variance** | | **Relative Efficiency** | |
|---|---|---|---|---|---|---|
| $\rho$ | $\pi_T$ | $\pi_S$ | Marginal | Spray | Empirical | Theoretical |
| 0.00 | 0.25 | 0.25 | 0.0018 | 0.0018 | 0.9998 | 1.0000 |
| 0.25 | 0.25 | 0.25 | 0.0018 | 0.0018 | 1.0140 | 1.0142 |
| 0.50 | 0.25 | 0.25 | 0.0018 | 0.0017 | 1.0607 | 1.0606 |
| 0.75 | 0.25 | 0.25 | 0.0018 | 0.0015 | 1.1545 | 1.1548 |
| 0.00 | 0.25 | 0.50 | 0.0009 | 0.0009 | 0.9998 | 1.0000 |
| 0.25 | 0.25 | 0.50 | 0.0009 | 0.0009 | 1.0106 | 1.0107 |
| 0.50 | 0.25 | 0.50 | 0.0009 | 0.0009 | 1.0475 | 1.0476 |
| 0.75 | 0.25 | 0.50 | 0.0009 | 0.0008 | 1.1305 | 1.1304 |
| 0.00 | 0.50 | 0.25 | 0.0013 | 0.0013 | 0.9997 | 1.0000 |
| 0.25 | 0.50 | 0.25 | 0.0013 | 0.0013 | 1.0214 | 1.0217 |
| 0.50 | 0.50 | 0.25 | 0.0013 | 0.0012 | 1.0996 | 1.1000 |
| 0.75 | 0.50 | 0.25 | 0.0013 | 0.0010 | 1.2999 | 1.3000 |

### 8 GTEx

#### 8.1 Detailed Methods

##### 8.1.1 Data Preparation

Genotypes and RNA transcript expression data were obtained from the NHGRI Genotype Tissue Expression (GTEx) Project, Version 7 (phs000424.v7.p2). The target outcome was gene expression in the substantia nigra (SSN; $n = 80$). Expression levels from three candidate surrogate tissues were considered:

1. Whole blood ($n = 369$), which has historically been used as a proxy tissue for eQTL analysis due to its accessibility [1].

2. Skeletal muscle ($n = 491$), which was the tissue with the largest number of genotyped subjects overall.

3. Cerebellum ($n = 154$), which was the subtype of brain tissue with the largest number of genotyped subjects.

For each analysis, the sample consisted of subjects with expression in either the SSN or the surrogate tissue (or both). Table **S10** partitions the number of subjects available for each analysis by missingness pattern, and figure **S10** depicts the squared Pearson correlation between gene expression in the target and surrogate tissues. Although we have not done so here, we note that a reasonable strategy for reducing the multiple testing burden while eliminating transcripts that are unlikely to benefit from surrogate analysis would be to require at least a minimum $r^2$ for inclusion of a transcript in the analysis.

Standard QC filters were applied to the genotype data using PLINK [2, 3]. SNPs were excluded from the analysis if the per-SNP missingness exceeded 10%. The analysis was restricted to common variants by requiring a minor allele frequency of at least 5%. Further simulation studies are needed to evaluate whether the Spray Wald test is applicable to rare variants; however, specialized methods are generally required for rare variant analyses [4]. To reduce multiple testing, variants in high linkage disequilibrium ($R^2 > 0.8$) were greedily pruned within a 1 Mb sliding window using PLINK's `clump` command. After filtering and pruning, 557,490 SNPs remained in the analysis. A transcript was considered *expressed* if the raw read count exceeded 5 for at least 20% of subjects in both the target and surrogate tissues. This definition was adopted to ensure that transcript expression was sufficiently strong to be distinguished from sequencing noise, and to eliminate from consideration transcripts that were unlikely to benefit from surrogate outcome analysis. Specifically, a transcript that is only expressed in one of the tissues cannot in principle benefit from surrogate outcome analysis because its cross-tissue correlation is zero. For sequences with multiple isoforms, only the isoform with the greatest variation in expression across subjects was retained.

**Supporting Table S10**: **Sample sizes for different target-surrogate outcome pairs.** In each case, the target outcome was gene expression in the substantia nigra. For retention in the analysis, a subject was required to have expression in at least one of the target or surrogate tissues, genotypes, and covariates.

| **Surrogate** | Complete Cases | Surrogate only | Target only | Overall |
|---|---|---|---|---|
| Blood | 47 (12%) | 322 (80%) | 33 (8%) | 402 |
| Muscle | 64 (13%) | 427 (84%) | 16 (3%) | 507 |
| Cerebellum | 66 (39%) | 88 (52%) | 14 (8%) | 168 |

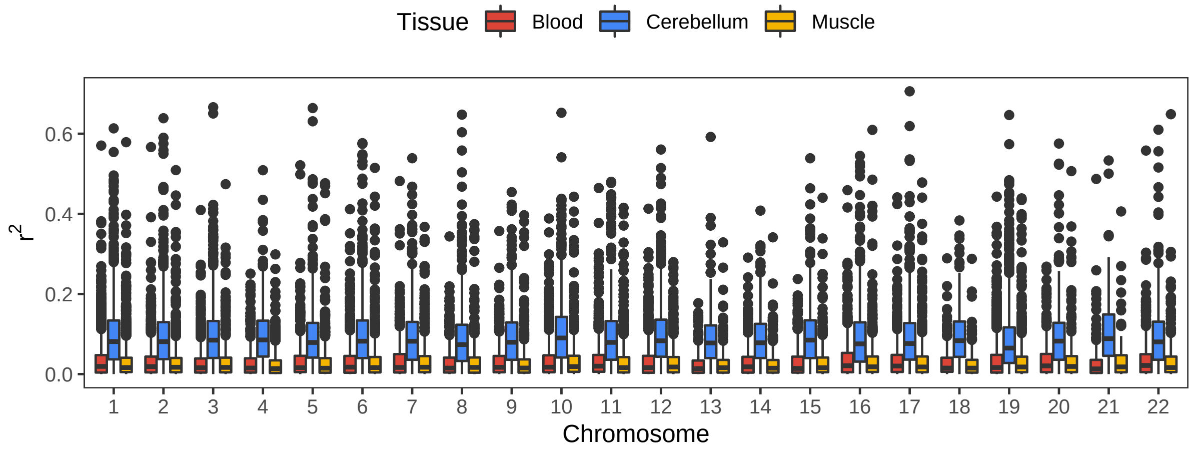

**Supporting Figure S10**: **Squared Pearson correlation between expression in the target and surrogate tissues.** In each case the target outcome is expression in SSN. Correlations for each *expressed* gene were calculated across complete cases.

##### 8.1.2 Association Testing

Association tests were performed for all variants in *cis* to a expressed transcript, defined as residing in the region spanning from 1 Mb upstream of the transcription start site to 1 Mb downstream of the transcription end site [3]. On median, there were 415 SNPs in *cis* to a transcript. A separate test of association was conducted for each *cis*-SNP by transcript pairing. Table (**S11**) presents the number of expressed genes and total number of SNP-transcript pairs for each surrogate tissue.

Target $T_i$ and surrogate $S_i$ expression levels were normalized within tissues via the rank-based inverse normal transformation (INT) [5], ensuring that the marginal distributions of each outcome were normal. All analyses included included age, sex, genotyping platform, and the top 3 genetic PCs as covariates. For the marginal analysis, transcript expression in the target issue (SSN) $T_i$ was regressed on genotype and covariates according to the association model:

$$T_i = \beta_0 + \beta_G g_i + \beta_A \text{Age}_i + \beta_S \text{Sex}_i + \beta_P \text{Platform}_i + \boldsymbol{\beta}'_{PC} \text{PCs}_i + \epsilon_{T,i}, \quad (1)$$

where $\epsilon_{T,i} \sim N(0, \Sigma_{TT})$. The target of inference was $\beta_G$. The null hypothesis $H_0 : \beta_G = 0$ was assessed using the standard Wald test, as implemented by base `R` [6]. Only subjects with observed expression in SSN were included in the marginal analysis. Note that expression in blood $S_i$ was not included in the marginal analysis because conditioning on $S_i$ in the regression model would change the interpretation of $\beta_G$ (see Section 5.1).

For the joint analysis, expression in SSN $T_i$ and expression in the surrogate tissue $S_i$ were jointly regressed on genotype and covariates according to the Spray model:

$$\begin{pmatrix} T_i \\ S_i \end{pmatrix} = \begin{pmatrix} \beta_0 + \beta_G g_i + \beta_A \text{Age}_i + \beta_S \text{Sex}_i + \beta_P \text{Platform}_i + \boldsymbol{\beta}'_{PC} \text{PCs}_i \\ \alpha_0 + \alpha_G g_i + \alpha_A \text{Age}_i + \alpha_S \text{Sex}_i + \alpha_P \text{Platform}_i + \boldsymbol{\alpha}'_{PC} \text{PCs}_i \end{pmatrix} + \begin{pmatrix} \epsilon_{T,i} \\ \epsilon_{S,i} \end{pmatrix},$$

where $(\epsilon_{T,i}, \epsilon_{S,i}) = \boldsymbol{\epsilon}_i \overset{\text{IID}}{\sim} N(\mathbf{0}, \boldsymbol{\Sigma})$. Estimation and inference were performed using the accompanying `R` package [7]. ECME iterations continued until the improvement in the observed data log likelihood dropped below $10^{-8}$. The null hypothesis $H_0 : \beta_G = 0$ was evaluated using the Spray Wald test. All subjects with expression in at least one of the target or surrogate tissues were included in the joint analysis.

**Supporting Table S11**: **Expressed transcripts and total SNP-transcript pairs by surrogate tissue.** In each case, the target outcome was gene expression in the substantia nigra. A transcript was considered expressed if the raw read count exceed 5 for 20% of subjects in both the target and surrogate tissues. The number of SNP-transcript pairs refers specifically to SNPs in *cis*.

| Surrogate | Expressed Transcripts | SNP-Transcript Pairs |
|---|---|---|
| Blood | 14,798 | 6,769,471 |
| Muscle | 15,292 | 6,953,630 |
| Cerebellum | 16,061 | 7,408,891 |

#### 8.2 Results

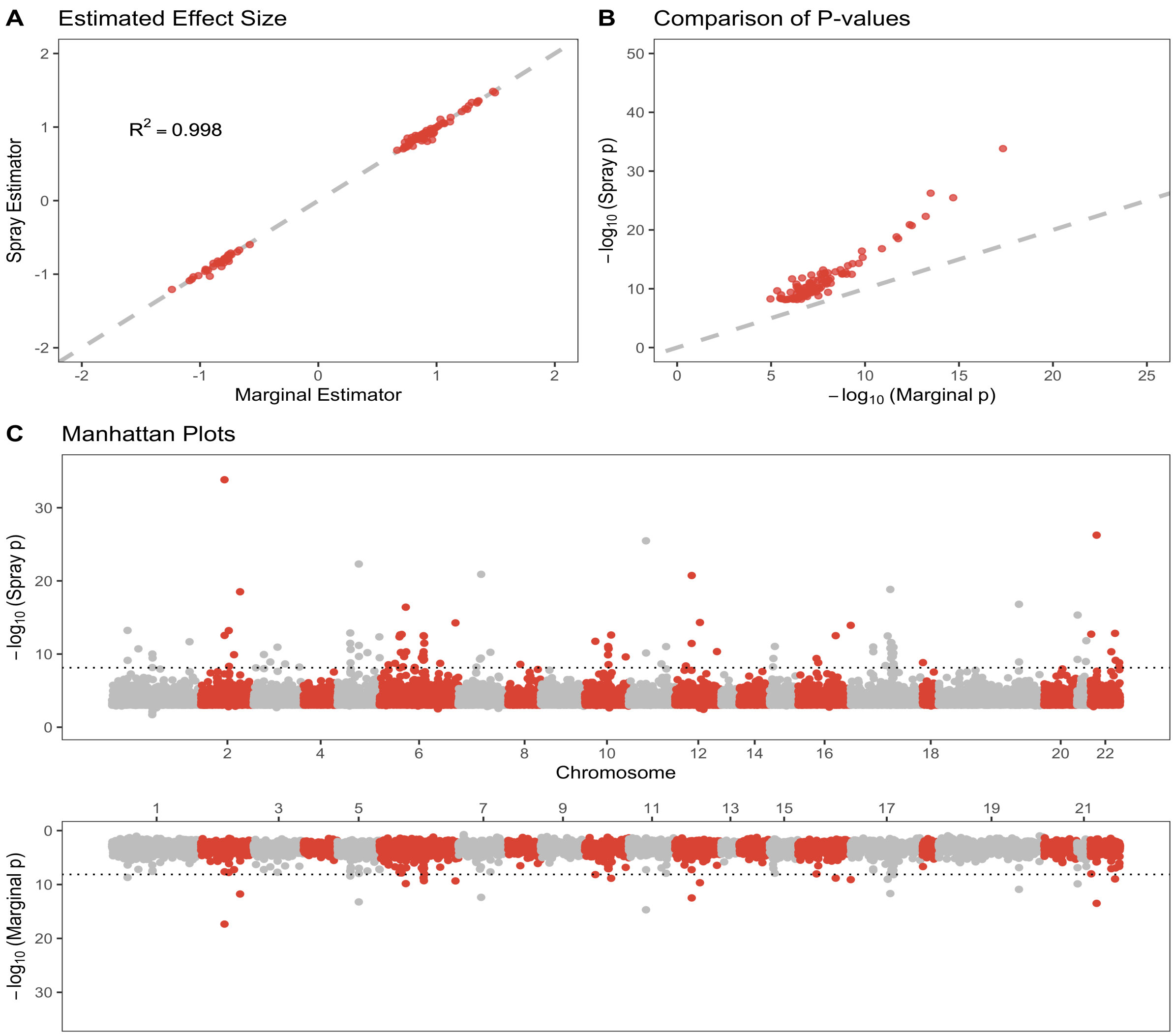

**Supporting Figure S11**: **Comparison of the marginal and joint (Spray) eQTL analyses of substantia nigra, using blood as the surrogate tissue.** A. Estimated effect size from the joint analysis vs. the estimated effect size from the marginal analysis for eQTL significant in at least one of the analyses. B. P-value from the joint analysis vs. p-value from the marginal analysis for eQTL significant in at least one of the analyses. C. Mirrored Manhattan plots comparing the p-values of the joint and marginal analyses by genomic position. Dotted line is the Bonferroni significance threshold.

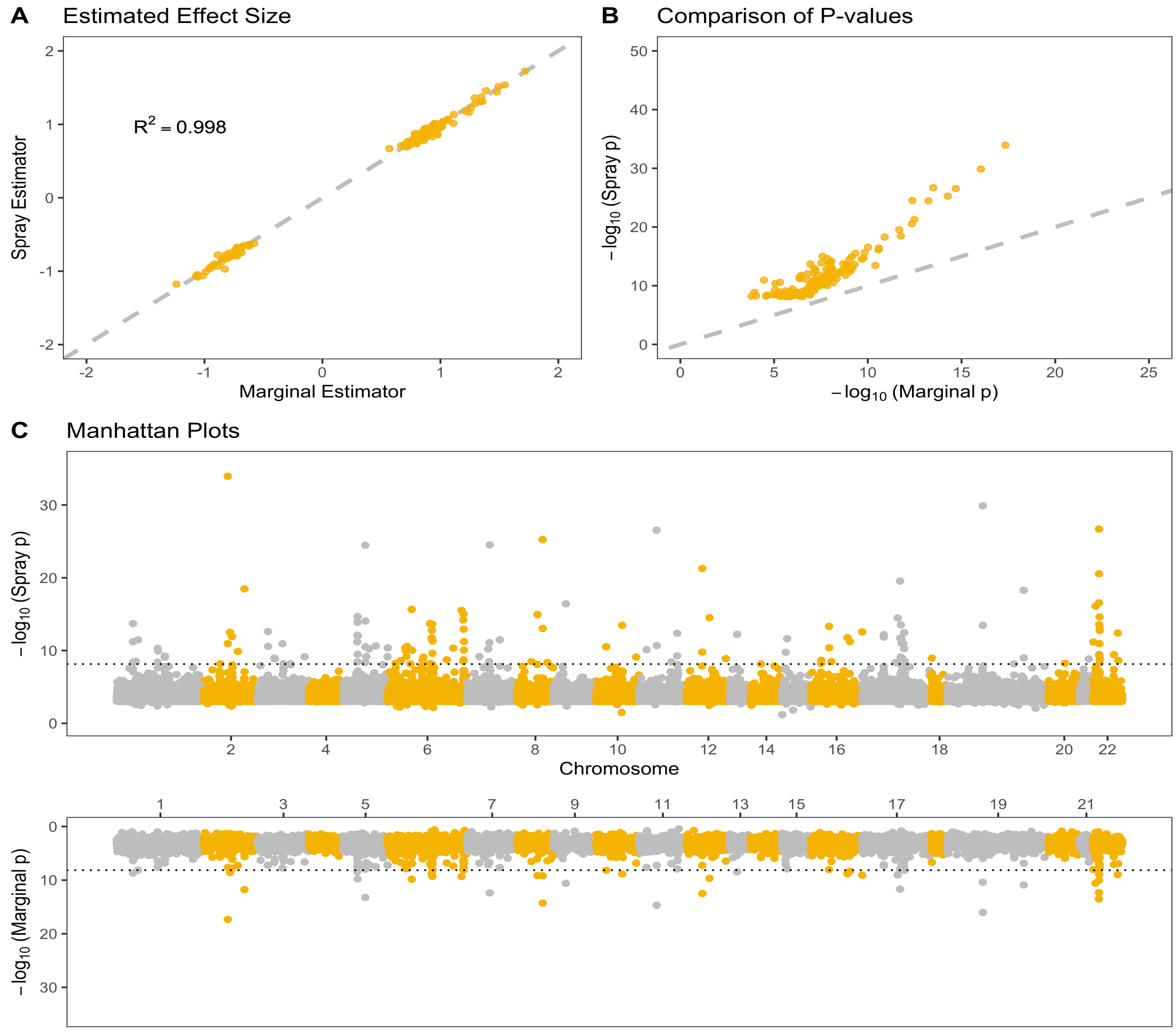

**Supporting Figure S12: Comparison of the marginal and joint (Spray) eQTL analyses of substantia nigra, using muscle as the surrogate tissue.** A. Estimated effect size from the joint analysis vs. the estimated effect size from the marginal analysis for eQTL significant in at least one of the analyses. B. P-value from the joint analysis vs. p-value from the marginal analysis for eQTL significant in at least one of the analyses. C. Mirrored Manhattan plots comparing the p-values of the joint and marginal analyses by genomic position. Dotted line is the Bonferroni significance threshold.
